# Engineering Large-Scale and Innervated Functional Human Gut for Transplantation

**DOI:** 10.1101/2025.05.22.655510

**Authors:** Holly M. Poling, Théo Noël, Akaljot Singh, Garrett W. Fisher, Konrad Thorner, Praneet Chaturvedi, Kalpana Nattamai, Kalpana Srivastava, Matthew R. Batie, Taylor Hausfeld, Amy L. Pitstick, Séverine Ménoret, Ignacio Anegon, Riccardo Barrile, Christopher N. Mayhew, Takanori Takebe, James M. Wells, Michael A. Helmrath, Maxime M. Mahe

## Abstract

A confined culture system (CCS) establishes methods to generate complex functional human gastrointestinal tissues. This approach yields large-scale innervated small intestinal, colonic and gastric organoids with an elongated tubular shape for both in vitro and in vivo studies. Transcriptomic and electrophysiological data demonstrate the co-development of a functional de novo enteric nervous system, which is absent from conventional organoids. When compared to traditional methods, CCS derived small intestinal, colonic and gastric organoids reached maturation supporting transplantation in half of the time, resulting in enhanced engraftment rates and sizes. Murine luminal content exposure within CCS organoids in vivo further augmented function, supporting the potential translational benefits required to model complex intestinal diseases. In summary, the CCS methodology simplifies current protocols while adding complexity and expediting the generation of clinically relevant functional gut tissues.

## Introduction

The ability to culture and manipulate human pluripotent stem cells (hPSC) has advanced the generation of specific tissues^1^. Translational embryology has enabled “organogenesis in a dish” in 3D environments, fostering more physiologically relevant models such as organoids^2,3^. Defined by their self-assembly and renewal, organoids mimic native tissue, include correct cell types, and replicate some organ functions^4^. These systems have been developed for many organs, including the gastrointestinal tract^5–10^, offering real-time visualization of organogenesis and bridging the gap between animal models and humans. hPSC-derived organoids hold immense potential for studying development, disease processes, cell-based therapies, and personalized drug discovery.

Despite these advantages, current organoid technologies face limitations in mimicking physiological environments in vitro, often requiring in vivo transplantation for continued tissue differentiation. Some hPSC-derived gastrointestinal tissues exhibit incomplete maturation, limiting their transplantation^11^. We hypothesized that methods to promote cellular complexities would be required to support both in vitro and in vivo maturation that will approximate native developing tissues^11,12^. hPSC-derived cells undergo normal maturation, thus initially have embryonic or fetal-like characteristics^12,13^. Methods to further develop these tissues are required to achieve the much-desired mature tissues to model human bowel. Using existing protocols, human intestinal organoids (HIOs) grow individually, reaching ∼1 mm after 28 days in culture, expanding to ∼1 cm with transplantation^14–16^. However, their spherical shape differs from the intestinal tube. We introduced strain into transplanted HIOs to induce maturation and enterogenesis, producing elongated tissues resembling the postnatal gut^12^. Recent methods to generate hindgut (HCO) and foregut (HGO, HaGO for antral) organoids have furthered understanding of gastrointestinal development and disease, though traditional culturing results in low engraftment rates and small sizes compared to HIOs^7,11^. Initial attempts to correct this were achieved in an assembloid approach by adding both splanchnic mesenchyme and neural crest cells^11^.

In this study, we focused on simplifying the methods required to achieve robust engraftment using a CCS. We designed a specialized scaffolding tray to provide a permissive environment for spheroid fusion (the 3D rounded structures that form from 2D monolayers upon differentiation, which are typically collected and replated for maturation into organoids). Loading a critical mass of spheroids into the scaffolding trays yielded longitudinal organoid structures. This device combined with 3D culture techniques to prime organoids successfully enhanced tissue growth and maturation, yielding organoids ten times larger than previous protocols. Notably, a de novo enteric nervous system (ENS) emerged, displaying selective neuronal excitation and contractile functions. Similar results were observed in colonic and gastric organoids. These structures readily engrafted, exhibited enhanced maturation, and developed a functional ENS. Here, we present a scalable organoid generation method emphasizing cellular confinement, advancing the production of complex organs for clinical applications.

## Results

### Implementation of a scaffolding tray for organoid generation

To impart a defined geometry upon intestinal organoids, a specialized scaffolding tray was designed to culture mid-hindgut spheroids and introduced into the traditional workflow (Figure 1a). The scaffolding tray was made by 3D-printing a mold with longitudinal grooves to culture a critical spheroid mass (Supplementary Figure 1a). Biocompatible polydimethylsiloxane (PDMS) was then cured within the mold and removed for use (Supplementary Figure 1b). This approach, termed the CCS, restricted spheroid growth within a defined space, while enhancing scalability. Typically, a single spheroid forms a single HIO, but here, approximately 4000 spheroids from spontaneous morphogenesis were used (Supplementary Figure 1c). They began as discrete spheroids at d0 that developed into a unified construct by d5 (Figure 1b). By d6, these structures were manually removed and replated in Matrigel for continued culture until d14 (Supplementary Figure 1d). When comparing small intestinal CCS (SI CCS) to conventional HIOs at d6 and d14, the geometry was maintained, and both simple columnar epithelium and supporting mesenchymal populations persisted (Figure 1c).

**Fig. 1.**
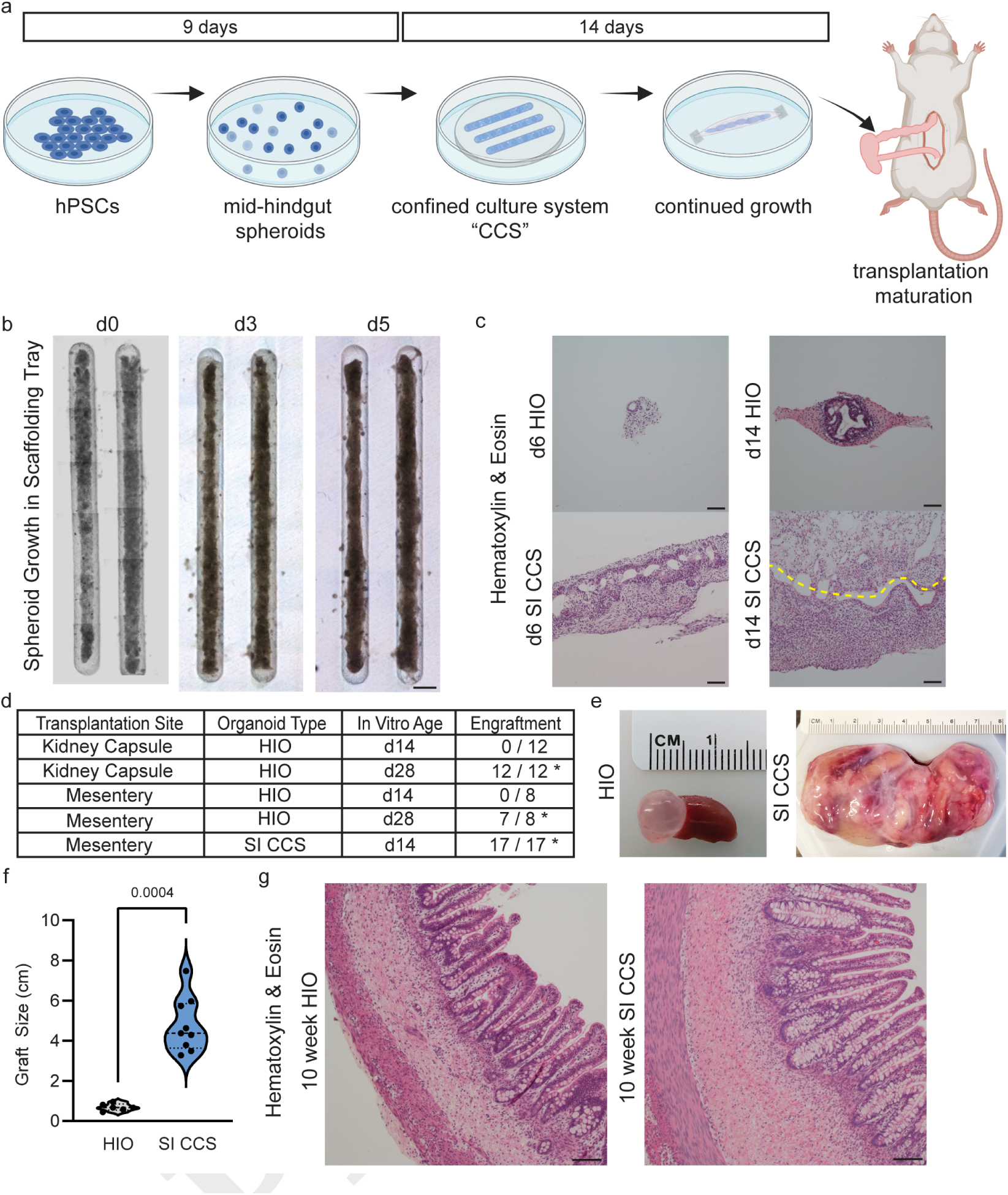
Employing a confinement strategy for the in vitro growth of small intestinal organoids. **a**, Experimental schematic. hPSCs were cultured and differentiated into mid-hindgut spheroids following published protocols, were then collected and seeded within the grooves of a specialized scaffolding tray, allowed to grow for six days and then removed from the tray and replated. Culture was continued until d14 at which time the elongated organoids were transplanted. **b**, Tile scan images of spheroids growing within the scaffolding tray grooves at d0, d3 and d5 of culture. Scale bar = 1 mm. **c**, Representative images of hematoxylin and eosin stained sections of conventionally generated HIOs and SI CCS at d6 and d14 of culture. Scale bar = 100 µm. **d**, Table of engraftment efficiencies for HIO and SI CCS. **e**, Representative gross images of HIO and SI CCS at the time of harvest (10 weeks post-transplantation). **f**, Violin plot of HIO and SI CCS graft sizes. **g**, Representative images of hematoxylin and eosin stained sections of conventionally generated HIOs and SI CCS 10 weeks after transplantation. Scale bar = 100 µm. *n ≥* 4 for histological staining.

### Reduction in Time to Engraftment and Enhanced Morphometric Features

After d14 in culture, SI CCS structures displayed robustness in size and cellular integrity, suggesting sufficient maturation for transplantation. This contrasts with previous findings requiring HIOs to be cultured for at least 28d before transplantation into the kidney capsule or mesentery of immunocompromised mice (Figure 1d, Supplementary Figure 1e)^14^. When transplanted at 14d, SI CCS achieved 100% engraftment (Figure 1d); however, the mouse’s abdominal space constrained growth, necessitating early tissue harvest. To overcome this, we used an immunocompromised rat model (*Rag1*and *Il2rg*-deficient rats, RRG) for all CCS transplants^17^. To assess whether host size influenced organoid growth potential, d28 HIOs were transplanted into mice and rats. Upon harvest, no significant size differences were observed, excluding host size as a variable. Previous studies also demonstrated transplantation site did not affect outcomes (Supplementary Figure 2a)^15^. Based on these findings, we refined animal use, transplanting conventional organoids into mice and CCS organoids into rats.

After 10 weeks, SI CCS and HIO grafts were harvested (Figure 1e, Supplementary Figure 2b). SI CCS grafts were significantly larger than HIOs, reaching widths of 8 cm (Figure 1f). Histological analysis confirmed a stereotypical intestinal phenotype (Figure 1g), with appropriate proliferative zonation as demonstrated by marker of proliferation Ki-67, MKI67, staining (Supplementary Figure 2c). Digestive enzyme Sucrase Isomaltase (SI) and functional Alkaline Phosphatase (ALP1) were demonstrated via immunofluorescence (Supplementary Figure 2d-e). Secretory epithelial cell types were also identified and observed through staining for Alcian Blue for mucin-producing goblet cells, chromogranin A (CHGA) marking enteroendocrine cells, and antimicrobial peptide alpha defensin 5 (DEFA5) labeling Paneth cells (Supplementary Figure 2f-h). Subepithelial telocytes, important supporting cells of the intestinal epithelium, were also observed and localized adjacent to the epithelium^16^ (Supplementary Figure 2i). Morphometric analysis revealed SI CCS had increased crypt depth, villus height, and muscle thickness compared to HIOs (Supplementary Figure 2j-l). No differences in crypt fission were observed, likely due to size plateauing under homeostatic conditions (Supplementary Figure 2m). Notably, SI CCS exhibited vast stretches of continuous epithelium spanning several centimeters (Supplementary Figure 2n), demonstrating enhanced epithelial architecture and structural maturity.

### Co-emergence of a de novo Enteric Nervous System

Looking beyond the epithelium, SI CCS exhibited a robust network of myenteric plexuses, appropriately localized between circular and longitudinal muscle layers (Figure 2a). Immunofluorescence staining confirmed neuronal presence using panneuronal marker Ubiquitin C-Terminal Hydrolase L1 (UCHL1, aka PGP9.5), and a human cellular marker, X-Ray Repair Cross Complementing 5 (XRCC5, aka KU80). This suggests, first, the presence of neurons, and second, that their origin was not due to host neuronal ingrowth (Figure 2b)^18^. As further confirmation, staining for additional pan-neuronal markers Tubulin Beta 3 Class III (TUBB3) and ELAV Like RNA Binding Proteins 3 and 4 (ELAV3/4, aka HUC/D) were also observed in SI CCS (Figure 2c). Excitatory cholinergic and inhibitory nitrergic neuronal subtypes, were also present in the SI CCS^19^. Cholinergic neurons were demonstrated through co-staining for Choline O-Acetyltransferase (CHAT) and Neurofilament Protein (NF200), while nitrergic neurons were shown through co-staining for Nitric Oxide Synthase 1 (NOS1) and NF200 (Figure 2d-e)^20^. Synaptic vesicles (Synapsin1, SYN1), and an intermediate filament protein nestin (NES), were also observed throughout the muscularis of SI CCS (Figure 2f)^20,21^. We also confirmed the presence of glial cells using S100 Calcium Binding Protein B (S100B) as a marker, which were closely associated with neurons (Figure 2g). Because interstitial cells of Cajal (ICCs) localize near neuronal bundles and serve as the pacemaker of the gut, we investigated their presence using KIT Proto-Oncogene, Receptor Tyrosine Kinase (KIT) and found that they were in close association with TUBB3+ cell bundles (Figure 2h).

**Fig. 2.**
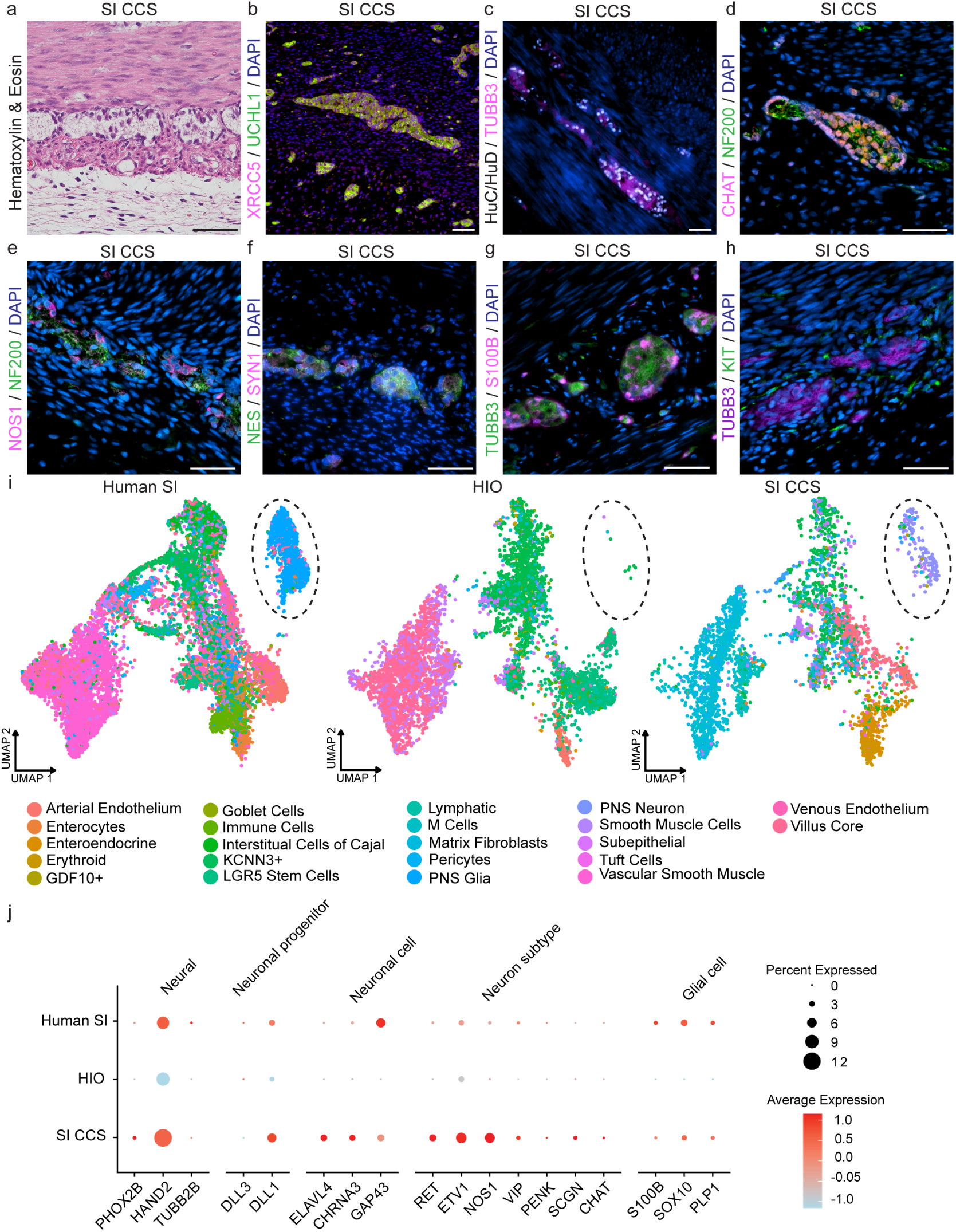
A robust ENS co-developed within SI CCS. **a**, Representative image of hematoxylin and eosin stained section of SI CCS 10 weeks after transplantation. **b**, Immunofluorescence for a human-specific marker (XRCC5, pink) and a pan-neuronal marker (UCHL1, green) in SI CCS. **c**, Immunofluorescence for two pan-neuronal markers (TUBB3, green; and HuC/HuD, white) in SI CCS. **d**, Immunofluorescence for neurofilaments (NF200, green) and a cholinergic neuronal marker (CHAT, pink) in SI CCS. **e**, Immunofluorescence for neurofilaments (NF200, green) and a nitrergic neuronal marker (NOS1, pink) in SI CCS. **f**, Immunofluorescence for intermediate neurofilaments (NES, green) and a synaptic vesicles (SYN1, pink) in SI CCS. **g**, Immunofluorescence for glia (S100B, pink) and neurons (TUBB3, green) in SI CCS. **h**, Immunofluorescence for neurons (TUBB3, purple) and ICCs (KIT, green) in SI CCS. **i**, UMAPs of snRNAseq datasets generated from the human small intestine, transplanted HIO and transplanted SI Groove with query transferred labels from curated reference atlas. Dashed oval indicates the localization of neuroglial cell clusters. **j**, Dot plot of transcription levels from snRNAseq datasets of genes related to the ENS. **b-h**, scale bars = 50 µm. *n ≥* 4 for all histological and immunofluorescence staining.

As an orthogonal method to confirm ENS presence, single nuclear isolations were performed on the human small intestine, transplanted HIOs, and transplanted SI CCS for single nuclear RNA sequencing (snRNAseq). A reference atlas was curated from three publicly available single cell RNA sequencing (scRNAseq) data sets, including STAR-FINDer^22^, the Gut Cell Survey^23^, and the Endodermal Organ Cell Atlas^24^. UMAP analysis confirmed successful integration of the datasets and reference (Supplementary Figure 3a). Label transfer analysis validated our protocol’s ability to isolate ENS cell types, as neurons and glia were identified in human samples. Transplanted HIOs lacked these populations, whereas SI CCS contained ENS-related populations, further confirming neuroglial identity (Figure 2i). We also defined the proportions of umbrella cell types (endothelium, mesenchyme, neural, immune, red blood cells, and epithelium) across sample types. We observed an increase in both mesenchymal and neuronal populations in SI CCS compared to HIO (Supplementary Figure 3b).

To specifically investigate the transcription levels of ENS-related genes, a dot plot was generated from the three data sets (Figure 2j). The average expression for genes related to neuronal progenitors, neuronal cells and subtypes, and glial cells was more robust in SI CCS than HIO which displayed little to no expression across the gene panel. The presence of neuronal subtype related gene expression led us to making neuronal subtype predictions utilizing the published datasets of Drokhlyansky et al., 2020 (Supplementary Figure 3c)^25^. Using this approach, putative excitatory motor neurons, putative inhibitory motor neurons and putative secromotor/vasodialator neurons were predicted within the SI CCS dataset further suggesting neuronal heterogeneity.

Our group previously generated HIOs with an ENS by adding exogenous neural crest cells to spheroids (HIO+ENS assembloids)^26^. To compare the ENS generated by both methodologies, we examined UCHL1+ myenteric plexuses in transplanted HIO+ENS, SI CCS, and human intestinal samples. The plexus size was significantly larger in SI CCS, suggesting that co-development within CCS enhances innervation more effectively than assembling ENS components separately (Supplementary Figure 3d-e).

### Neuromuscular Function in SI CCS

Organ bath chamber assays were conducted to assess neuromuscular function in SI CCS. Continuous monitoring of contractile activity revealed rhythmic isometric force contractions after equilibration (Figure 3a). To contextualize this non stimulated activity, transplanted HIO+ENS and human small intestinal samples were also analyzed. Contractile amplitude was significantly greater in SI CCS than in HIO+ENS and comparable to human intestine (Figure 3b). Selective nerve excitation through electric field stimulation (EFS) was then performed on all three sample types (Figure 3c-e). HIO+ENS showed minimal contractile activity, consistent with prior findings^26^, while SI CCS and human intestine exhibited strong contractile responses, confirming successful nerve excitation. ENS dependence was validated by tetrodotoxin (TTX) application, which significantly reduced contractile responses upon repeated EFS (Figure 3f). To investigate the functional contributions of cholinergic and nitrergic neurons, which were demonstrated through protein expression, nitric oxide synthetase (nNOS)-expressing neurons were inhibited with NG-nitro-l-arginine methyl ester (L-NAME), followed by cholinergic inhibition using atropine^26^. Contractile activity associated with EFS was measured before and after each treatment (Figure 3g). These data demonstrated that SI CCS contractions involved both nitrergic and cholinergic neuronal activity, with the cholinergic component being more dominant (Figure 3h).

**Fig. 3.**
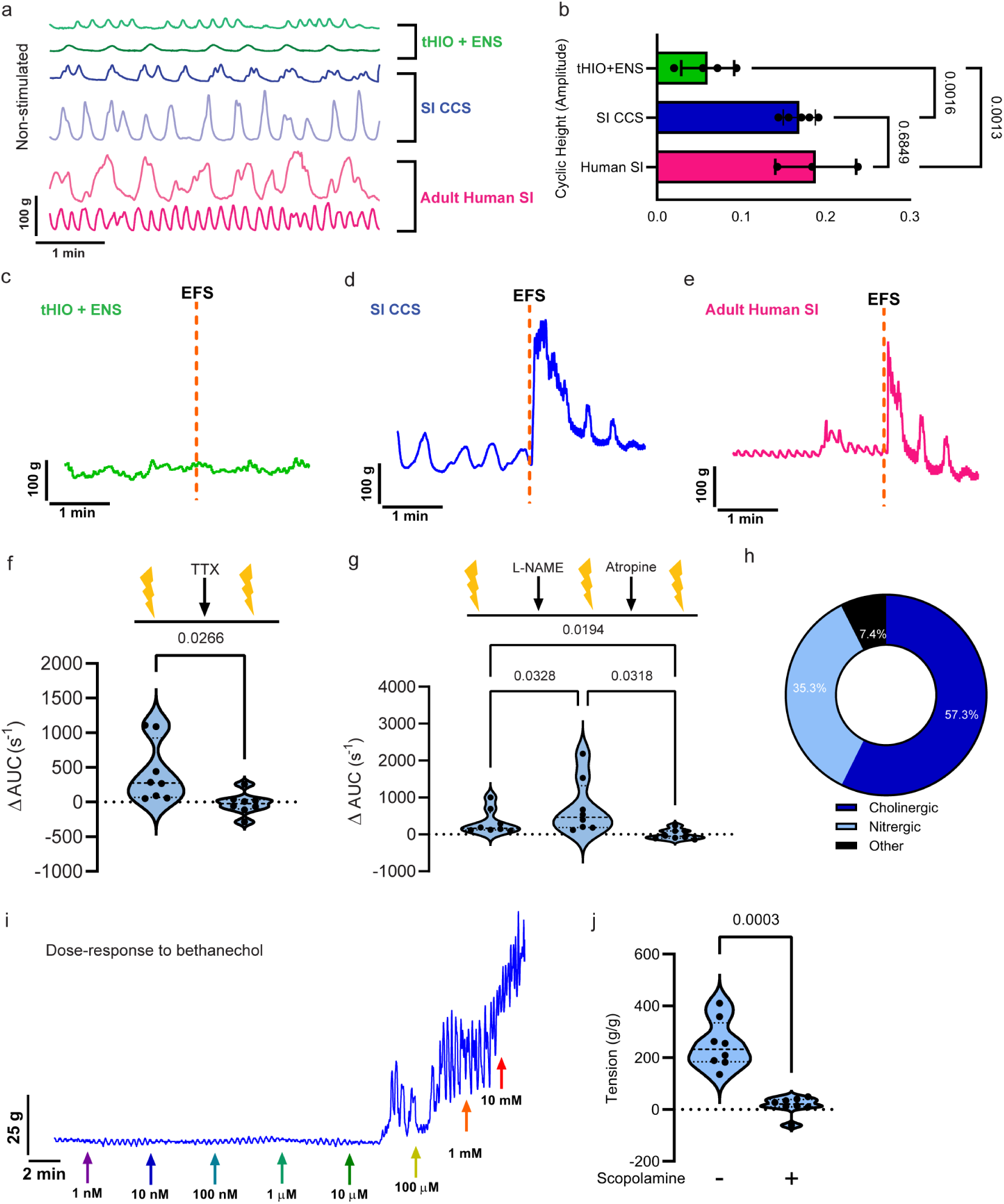
SI CCS contained a functional ENS that regulates smooth muscle contractility. **a**, Isometric force contractions of muscle strips from tHIO+ENS, SI CCS, and Adult Human SI after an equilibration period. **b**, Bar graph (mean±SD) of the average cyclic height of contractile tracings from tHIO+ENS, SI CCS, and Adult Human SI. SI CCS and Adult Human SI values demonstrated no significant difference between groups. **c**, Contractile activity tracing in tHIO+ENS with EFS stimulation (dashed line indicates the administration time). **d**, Contractile activity tracing in SI CCS with EFS stimulation (dashed line indicates the administration time). **e**, Contractile activity tracing in Adult Human SI with EFS stimulation (dashed line indicates the administration time). **f**, TTX inhibition of ENS activity in SI CCS. Violin plot of the change in the area under the curve before and after TTX administration; after TTX neuronal response was significantly reduced. **g**, Nitrergic and cholinergic inhibition of ENS activation. Violin plot of the change in the area under the curve following control EFS stimulation, followed by L-NAME treatment, another EFS stimulation, followed by Atropine treatment, and a final EFS stimulation in SI CCS. Each EFS response was significantly different from one another. **h**, Average calculated neuronal subtype functional contribution from experiments in panel g. **i**, Representative bethanechol dose-response tracing of contractile activity in a transplanted SI CCS; with dosage increases, tensile activity increased. Colored arrows indicate the timing of logarithmic bethanechol doses. **j**, Calculated maximal and minimal tissue tension of SI CCS; scopolamine was used to induce muscle relaxation. n=8 for organ bath experiments.

Smooth muscle function, independent of ENS influence, was assessed by administering bethanechol, a muscarinic receptor agonist. Contractile force increased in a dose-dependent manner (Figure 3i), confirming functional smooth muscle formation in SI CCS. These effects were reversible, as smooth muscle relaxation was induced with scopolamine, a muscarinic antagonist (Figure 3j). Together, these results demonstrate that SI CCS successfully generates functional smooth muscle, which operates independently and in conjunction with ENS stimulation.

### The CCS Promotes in vitro ENS Development

Previously, generating organoid tissues with an ENS required adding exogenous neural crest cells, differentiated separately and incorporated with spheroids^11,26,27^. Surprisingly, SI CCS developed a functional ENS without this method. To gain insight into the emergence of the ENS in SI CCS, transcriptomic analysis was performed at in vitro d14, first between HIO and SI CCS. When examining the heatmap of the two groups, d14 HIOs and d14 SI CCS segregated and presented with transcriptional differences (Supplementary Figure 4a). Differentially expressed genes in SI CCS highlighted enriched gene ontologies related to ENS development (Supplementary Figure 4b). The top five biological processes and cellular components were all linked to early ENS formation, suggesting that CCS promotes endogenous ENS development within the organoids.

### The in vitro SI CCS ENS Represents an Early Developmental State

Next, we compared the de novo ENS in SI CCS to that in HIO+ENS. EFS failed to induce responses in engrafted HIO+ENS, but produced robust responses in SI CCS, indicating functional differences. To understand factors that may have contributed to this functional difference related to engraftment, single cell RNA sequencing (scRNAseq) was performed on d28 HIO+ENS and d14 SI CCS, their respective stages for transplantation. First, a harmonized UMAP was generated and four umbrella cell types (endothelial, epithelial, mesenchymal and neural) were identified (Figure 4a). Then, the integration of the two sample types was explored through another UMAP, color-coded by group (Figure 4a). A bar graph was generated using the four umbrella cell types to examine differences in cell type proportions (Figure 4b). Associated raw event counts were also calculated (Supplementary Figure 4c). Interestingly, the d14 SI CCS had a smaller neural proportion than the d28 HIO+ENS. However, for the other categories (endothelial, epithelial, and mesenchymal), d14 SI CCS demonstrated a larger proportion than d28 HIO+ENS. This may be an artifact of dissociation, or because the neural population was exogenously provided within the HIO+ENS. Irrespective of the cause, we wanted to interrogate further the neural populations captured in both groups. As a first line assessment, feature plots visualized key ENS-related genes (Figure 4c). *SRY-Box Transcription Factor 10* (*SOX10*), a transcription factor that helps maintain neural cells in an undifferentiated proliferative state, was rarely observed within the d28 HIO+ENS, but well represented within d14 SI CCS^28^. *Forkhead Box D3* (*FOXD3*), a transcription factor expressed in early neural crest cells, which is pivotal in maintaining their multipotency, demonstrated a similar expression pattern^29^. *ELAV Like RNA Binding Protein 4* (*ELAVL4*), a marker for cells that have committed to a neuronal fate was more broadly expressed within the neuronal cluster of d28 HIO+ENS than in d14 SI CCS^30^. *Endothelin Receptor Type B* (*EDNRB*), which plays an early role in neural migration and the prevention of premature differentiation, seemed represented to a similar level in the neural clusters of both d28 HIO+ENS and d14 SI CCS^31,32^.

**Fig. 4.**
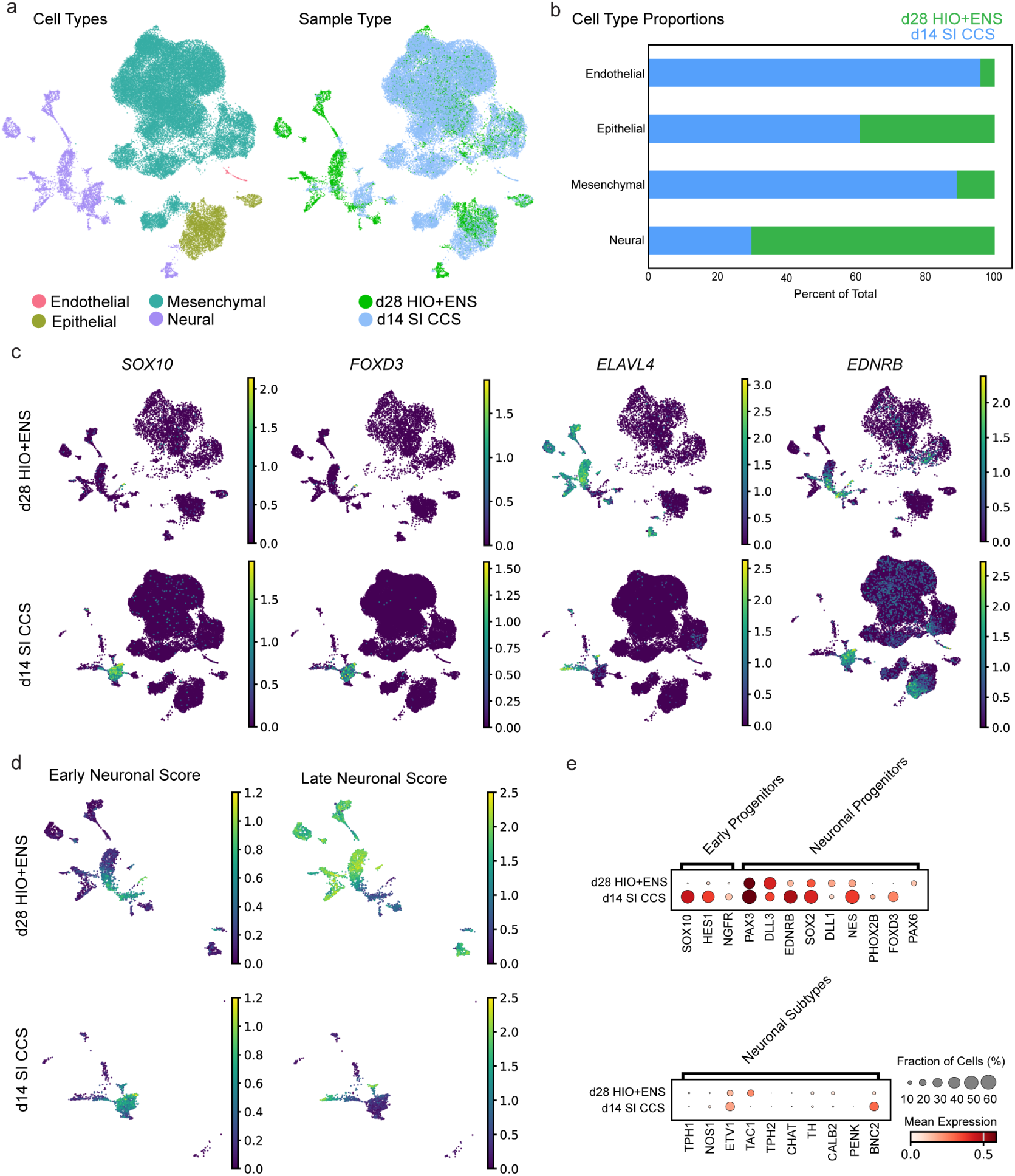
Transcriptional profiling reveals an enrichment in early ENS progenitors within SI CCS pre-transplantation. **a**, UMAPs of in vitro d28 HIO+ENS and d14 SI CCS scRNAseq color coded by cell type identity (grouped as Endothelial, Epithelial, Mesenchymal and Neural), and according to sample type. **b**, Bar graph of cell type proportions in d28 HIO+ENS and d14 SI CCS. **c**, Feature plots highlighting transcription of genes SOX10, FOXD3, ELAVL4, and EDNRB in d28 HIO+ENS and d14 SI CCS. **d**, UMAPs of the isolated neural cluster in d28 HIO+ENS and d14 SI CCS indicating early and late neuronal scores across each sample type. d28 HIO+ENS demonstrate more widespread late neuronal scores, while d14 SI CCS demonstrate early neuronal scores. **e**, Dot plot of transcription levels from scRNAseq datasets of genes related to the early progenitors, neuronal progenitors and neuronal subtypes.

To further assess neuronal maturity, early and late neuronal scores were established based on well-characterized ENS differentiation markers. d28 HIO+ENS exhibited high late neuronal scores, whereas d14 SI CCS demonstrated high early neuronal scores (Figure 4d). Neural subpopulations were then stratified using the Fetal Gut Cell Atlas enteric dataset (Supplementary Figure 4d). Best-match predictions revealed that early progenitor subclusters, including enteric neural crest cells (ENCC)/glia progenitors, cycling ENCC/glia, neuroblasts, and cycling neuroblasts, accounted for ∼40% of the neural cluster in d28 HIO+ENS but over 80% in d14 SI CCS (Supplementary Figure 4e). Dot plots confirmed the enrichment of early progenitor markers in d14 SI CCS compared to d28 HIO+ENS (Figure 4e). These findings indicate that the ENS within SI CCS is at an earlier developmental stage, with a higher proportion of progenitors, potentially contributing to its enhanced functional capacity upon engraftment.

### Extending CCS Technique to Generate Additional Gastrointestinal Tissue Types

Current gastrointestinal organoid protocols often result in poor engraftment and small size, limiting physiological studies and clinical applications^5–7,9,11^. Assembloid approaches, differentiation of key components separately before recombination, can mitigate these issues but also introduces new challenges related to cell ratios and timing^11,25,26^. To address this, we applied our CCS methodology to generate colonic and gastric tissue, streamlining organoid development and improving overall transplant potential.

We explored whether the CCS methodology could generate colonic and gastric tissue, given those protocols also rely on spheroid morphogenesis from monolayers. Using a previously published protocol, we introduced transient BMP activation for organoid posteriorization^7^. Similar to SI CCS, Colonic CCS (C CCS) successfully engrafted at d14, whereas conventionally generated HCOs required an additional 14d of maturation (Figure 5a). Engraftment efficiency was significantly improved with the scaffolding tray (Figure 5a). After ten weeks, HCOs and C CCS were harvested, with C CCS reaching widths of ∼6 cm (Figure 5b-c). Foregut spheroids^6,11^ placed in scaffolding trays also improved antral gastric tissue (G CCS) engraftment and size (Figure 5d-f).

**Fig. 5.**
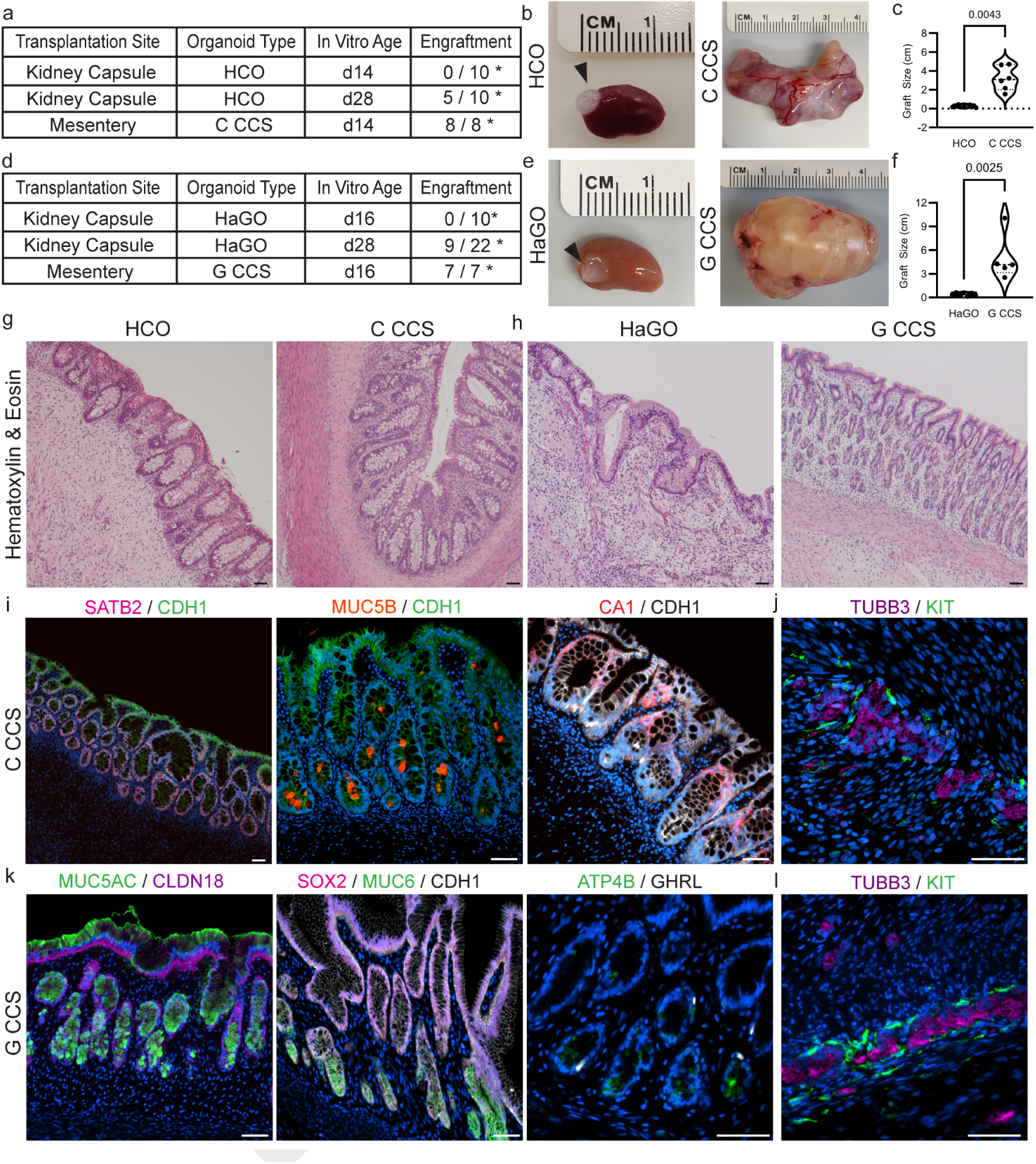
The scaffolding tray methodology successfully generated colonic and gastric tissue. **a**, Engraftment rates of conventionally generated HCOs and C CCS. **b**, representative gross images of HCO and C CCS at the time of harvest (10 weeks post-transplantation). **c**, Violin plot of HCO and C Groove graft sizes. **d**, Engraftment rates of conventionally generated HaGOs and G CCS. **e**, Representative gross images of HaGO and G CCS at the time of harvest (10 weeks post-transplantation). **f**, Violin plot of HaGO and G Groove graft sizes. **g**, Representative images of hematoxylin and eosin stained sections of conventionally generated HCO and C CCS 10 weeks after transplantation. **h**, Representative images of hematoxylin and eosin stained sections of conventionally generated HaGO and G CCS 10 weeks after transplantation. **i**, Representative immunofluorescence staining for distal and colonic markers in transplanted C CCS: distal transcription factor (SATB2, pink) and epithelium (CDH1, green), a colonic mucin (MUC5B, orange) and epithelium (CDH1, green), and a colonic epithelial enzyme (CA1, red) and epithelium (CDH1, white). **j**, Representative immunofluorescence for neurons (TUBB3, purple) and ICCs (KIT, green) in transplanted C Groove. **k**, Representative immunofluorescence staining for gastric markers in transplanted G CCS: gastric mucin (MUC5AC, green) and gastric epithelium (CLDN18, purple), a transcription factor (SOX2, purple), an antral gastric mucin (MUC6, green) and epithelium (CDH1, white), and a parietal cell transporter (ATP4B, green) and gastric hormone ghrelin (GHRL, white). **l**, Representative immunofluorescence for neurons (TUBB3, purple) and ICCs (KIT, green) in transplanted G Groove. **g-l**, scale bars = 50 µm. *n ≥* 4 for all histological and immunofluorescence staining.

Histologically, C CCS displayed a characteristic colonic phenotype lacking villi with a well-defined muscularis, which was largely absent in transplanted HCOs (Figure 5g). G CCS exhibited organized pits and glands, however more a rudimentary structure was found in conventional HaGO (Figure 5h). To further support the tissue type identity of C CCS and G CCS organoids, immunofluorescence for regional markers was performed. C CCS were found to express SATB Homeobox 2 (SATB2), a marker of distal gut epithelium^7^, Mucin 5B (MUC5B), a gel-forming mucin expressed by a subset of colonic Goblet cells^33^, as well as a colonic enzyme carbonic anhydrase 1 (CA1)^34^ supporting that the colonic patterning was successful (Figure 5i). Next, pan-neuronal staining was performed to determine if neuronal codevelopment persisted with C CCS generation. Interestingly, nerves were again observed in close association with ICCs (Figure 5j). G CCS were found to express Mucins 5AC and 6 (MUC5AC, MUC6)^33^, gel forming mucins expressed by some gastric Goblet cells, Claudin 18 (CLDN18)^35^, a gastric transmembrane protein, and SRY-Box Transcription Factor 2 (SOX2)^36^, expressed during stomach development (Figure 5k). Parietal cells, marked by ATPase H+/K+ Transporting Subunit Beta (ATP4B) expression^37^, as well as the gastric hormone ghrelin (GHRL)^38^ were also observed within G CCS samples, supporting their successful gastric patterning (Figure 5k). Then, neuronal co-development was demonstrated through pan-neuronal staining in G CCS samples, establishing that the CCS fostered the co-emergence of an ENS in all three contexts investigated (Figure 5j).

Next, we investigated if the ENS in the C CCS and G CCS tissues was functional through organ bath chamber assays like those performed on SI CCS. First, when interrogating the C CCS, rhythmic isometric force contractions were observed after a period of equilibration (Supplementary Figure 5a). Next, we investigated if mature neuromuscular coupling was also present through selective nerve excitation. EFS pulses were administered, and contractile activity was observed, indicating that the ENS regulated smooth muscle (Supplementary Figure 5b). To confirm that this phenomenon was ENS dependent, neuronal activity was inhibited with TTX, yielding a significantly reduced contractile response after another EFS application (Supplementary Figure 5c). Smooth muscle function was interrogated by the administration of bethanechol followed by scopolamine, in which contraction and relaxation were successfully induced (Supplementary Figure 6d-e). These assessments were repeated on G CCS, yielding similar results (Supplementary Figure 5f-j). Thus, CCS-based methodology successfully generates colonic and gastric tissue, both featuring a functional ENS.

### Proof of Concept for the Clinical Applicability of SI CCS

Exposing intestinal organoids to luminal content in vivo remains a challenge. Our group previously developed the Tie In procedure, where an engrafted organoid was anastomosed to the host’s bowel, enabling the study of intestinal adaptation from a sterile to a fed state with bacterial colonization and nutrient exposure (Supplementary Figure 6a)^39^. However, high mortality limited long-term studies (Supplementary Figure 6b). However, by 5d we observed the establishment of continuous epithelia between the host and transplanted organoid as demonstrated through immunofluorescence staining for CDH1 and XRCC5 (Supplementary Figure 6c). The stereotypical intestinal proliferative zonation of the host and organoid remained intact after the procedure as observed by immunofluorescence staining for MKI67 (Supplementary Figure 6c). Additionally, histological staining for bacteria was performed and multiple bacterium phenotypes were observed near the organoid’s epithelium (Supplementary Figure 6d). Organoid luminal contents were cultured, confirming bacterial outgrowth exclusively in the Tie In group (Supplementary Figure 6e). In response to luminal exposure, increased mucin production was observed histologically at d5, with elevated acidic and strongly sulfated mucosubstances (Supplementary Figure 6f). By d12, mucin levels declined but remained higher than in controls.

Transepithelial resistance (TEER) and paracellular permeability were also evaluated through electrophysiology at the d5 timepoint and it was similar between groups, however short circuit current (Isc), a measure of active ion exchange, was reduced in Tie Ins suggesting an increase in negative ion trafficking (Supplementary Figure 6g). Despite these insights, the model had substantial challenges, including the need for highly skilled surgeons, strict dietary control, and small organoid size limiting downstream analysis. To overcome these, we adapted the Tie In procedure using SI CCS (Figure 6a).

**Fig. 6.**
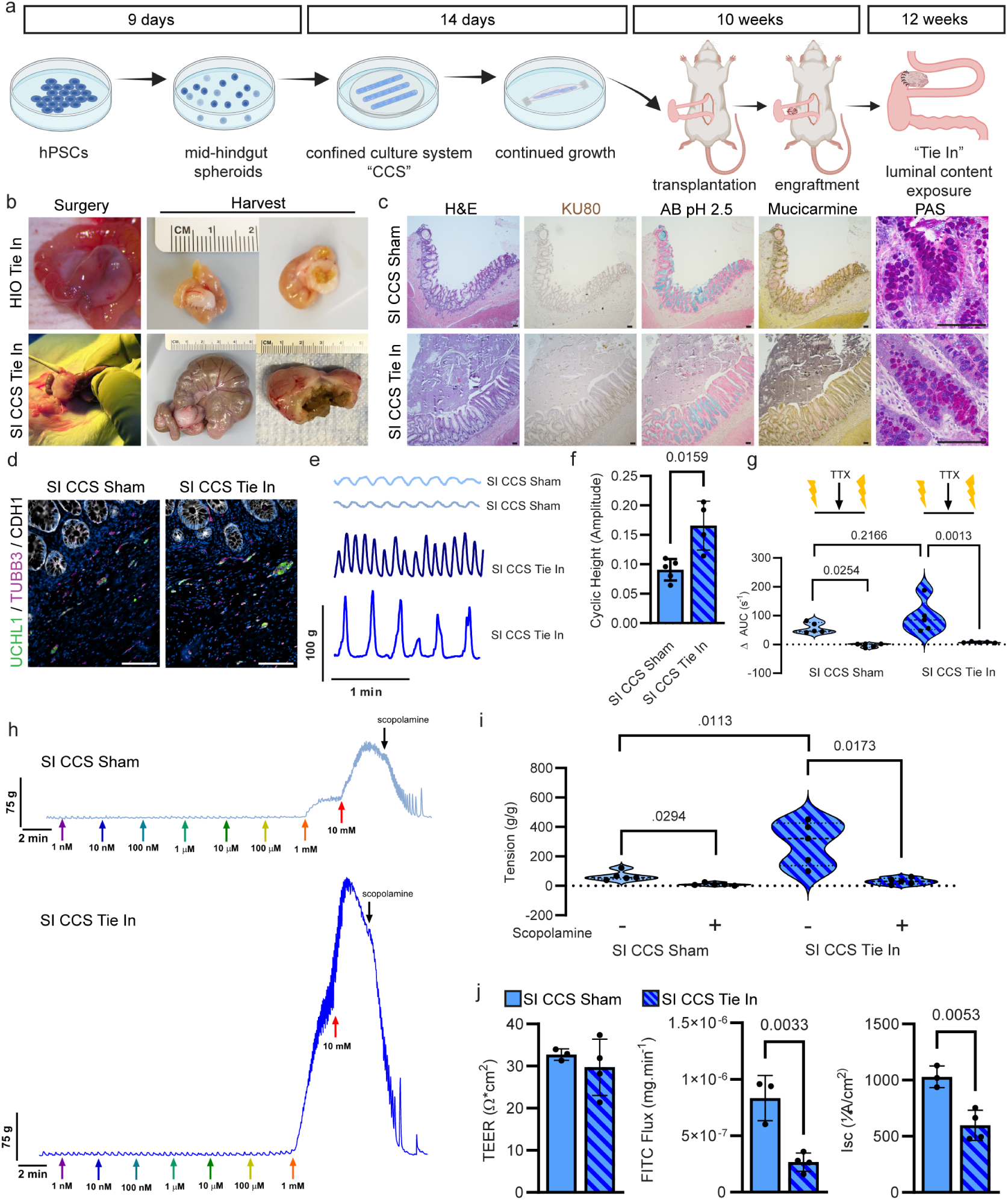
SI CCS robustly modeled adaptation to the fed intestinal state. **a**, Experimental schematic describing the generation and transplantation of SI CCS, engraftment, and secondary “Tie In” procedure to expose the engrafted tissue to host luminal content. **b**, Representative photographs of mouse HIO Tie In and rat SI CCS Tie In at the time of surgery and harvest. **c**, Representative staining panels of mucin profiles on serial sections of SI CCS Sham and SI CCS Tie In. Violin plot of HfGO and G Groove graft sizes. **d**, Representative immunofluorescence images of SI CCS Sham and SI CCS Tie In stained for neurons (UCHL1, green and TUBB3, purple) and epithelium (CDH1, white). **e**, Isometric force contractions of muscle strips from SI CCS Sham and SI CCS Tie In after an equilibration period. **f**, Bar graph (mean±SD) of the average cyclic height of contractile tracings from SI CCS Sham and SI CCS Tie In. **g**, TTX inhibition of ENS activity in SI CCS Sham and SI CCS Tie In. Violin plot of the change in the area under the curve before and after TTX administration; after TTX neuronal response was significantly reduced for both groups. **h**, Contractile activity tracings in SI CCS Sham and SI CCS Tie In in response to logarithmic doses of bethanechol, followed by scopolamine administration. Arrowheads indicate administration timings. **i**, Calculated maximal and minimal tissue tension of SI CCS Sham and SI CCS Tie In. **j**, Bar graphs (mean±SD) of functional epithelial characteristics in SI CCS Sham and SI CCS Tie In as measured in an Ussing Chamber assay. From left to right: transepithelial resistance, paracellular permeability and short circuit current. All scale bars = 100 µm. n=5 for organ bath experiments and n=3 (Sham) or 4 (Tie In) for Ussing chamber experiments.

The CCS model in rats significantly reduced mortality (Supplementary Figure 6c) and allowed for the dietary transition back to chow, enabling extended 12 week studies. Larger organoid tissue facilitated more comprehensive histological and electrophysiological analyses, while improved survival and engraftment refined animal usage (Figure 6b). For these procedures, part of the SI CCS was transplanted into the host’s mesentery to facilitate subsequent anastomosis to the cecum, chosen for its high microbial density and permissive anatomy. Like the mouse model, luminal contents filled the SI CCS Tie In at harvest (Figure 6b), and histology confirmed adaptive architecture with visible luminal contents (Figure 6c). Mucin profiles in SI CCS Tie Ins were homeostatic and heterogeneous, consistent with d12 HIO Tie Ins (Figure 6c). Next, we characterized functional maturation within the system. Interestingly, staining for pan-neuronal markers (TUBB3 and UCHL1) revealed continued ENS development within the SI CCS Tie In and sham (Figure 6d). At ten weeks, most of the neuronal structures were associated with the smooth muscle layers. However, at the 22-week time point, a more consistent incorporation was observed throughout the full thickness of the tissue, particularly in the submucosal space, in both groups. This indicated that the ENS development had not yet plateaued at the ten-week time point and that neuronal expansion occurred and was independent of the tie in the procedure. During adolescence, substantial architectural remodeling and changes occur within the ENS, particularly in the submucosal regions. This correlates to the most notable changes observed in the SI CCS tissues between 10 and 22 weeks of transplantation^40^.

When examining contractile tracings of the SI CCS Tie In and sham under non-stimulated conditions, the SI CCS Tie In had significantly increased amplitudes compared to sham (Figure 6e-f). EFS responses were comparable in both groups (Figure 6g), with contractions significantly reduced after tetrodotoxin (TTX) treatment, confirming ENS dependence (Figure 6g). These data suggest that luminal content exposure enhances ENS maturation beyond the effects of engraftment duration. Next, muscle tone was explored between the groups by administering logarithmic doses of bethanechol. During this experiment, live tracings of the contractile activity demonstrated a more robust response from SI CCS Tie In than sham (Figure 6h). The calculated maximum tension was significantly increased in SI CCS tie-in compared to the sham, while relaxation was successfully induced at a similar level in both groups (Figure 6i).

When evaluating epithelial function, TEER showed no significant differences between SI CCS Tie In and sham, but paracellular permeability was significantly reduced, indicating improved barrier function related to luminal content exposure (Figure 6j). Consistent with early d5 HIO Tie-In results, Isc was reduced in SI CCS Tie In (Figure 6j, Supplementary Figure 6g). This line of experimentation demonstrates the utility of the CCS platform for vigorous developmental modeling and manipulation. Together, these mouse and rat-derived data suggest that transplanted organoids undergo substantial adaptation processes upon luminal content exposure and that the organoid tissues thrive and mature in physiological conditions.

## Discussion

Human gastrointestinal organoids serve as valuable models for studying development and disease. Cultured from hPSCs, these 3D systems closely resemble native tissues and have advantages to animal models. Numerous groups have been using organoids and introducing ways to improve them for basic science and translational studies^11,26,27,41–45^. The field of tissue engineering and organoid technologies is rapidly co-evolving with increased diversity in the organoids/organs that can be generated and their level of complexity^6,9,11,26,44,46–48^. Personalized medicine applications have been employed and strides toward clinical translation are underway^43,49,50^.

With this study, we have leveraged the ease of soft lithography to manufacture scaffolds that were incorporated into otherwise traditional organoid culture conditions to generate an improved organoid for transplantation. The simplicity of this experimental approach makes this methodology widely accessible and straightforward to execute. We have established a more reliable and scalable method for generating transplantable gastrointestinal tissues using this technique— some reaching eight centimeters with clinically relevant epithelial surface areas^51^. Recently, Takahashi et al. developed centimeter-scale HIOs using a rotating bioreactor, requiring an assembloid approach to combine separately generated spheroids and mesenchyme^45^. While both methods enabled elongated structures through collective behavior, the CCS uniquely supported de novo ENS development^45^. Additionally, CCS-driven maturation accelerated in vivo engraftment by two weeks compared to existing protocols, with enhanced efficiency—an e ssential factor for t ranslating o rganoid technologies into clinical applications^7,11,14,26,52,53^.

Notably, CCS organoids developed a functional ENS without an assembloid approach, the addition of separately differentiated neuroglial cells, representing a paradigm shift in gastrointestinal tissue engineering. The ENS governs key functions, including motility, fluid regulation, and blood flow, making its replication critical for physiological modeling. While traditionally attributed to migratory neural crest cells, there is recent data in support of the additional origins of the ENS^54–58^. While the CCS-based ENS origins remain unidentified, this model offers a unique system to investigate t his. Especially, given that the CCS methodology gave rise to neuronal diversity that was previously unachieved in the assembloid HIO+ENS system^26^. Both inhibitory and excitatory neuronal subtypes were identified using snRNA-seq, protein expression analysis, and electrophysiological assays. This platform could aid in understanding ENS-related diseases, such as Hirschsprung’s disease, characterized by the absence of functional ganglion cells, and may provide a neuroglial cell source for conditions involving developmental deficits^59^.

Clinical translation of organoid-based therapies requires demonstrating functional integration within physiological environments. As proof of concept, we utilized a Tie In model to confirm t hat CCS-derived i ntestinal t issues can successfully integrate with host gut architecture and function when exposed to luminal contents. This demonstrated organoids not only engraft but also adapt to host physiology, mimicking postnatal gut maturation. Barrier integrity, epithelial transport, and neuromuscular activity were all maintained or enhanced, supporting their therapeutic potential for intestinal failure. Additionally, the extended post-operative timeframe of the Tie In model allowed ENS components to migrate into the submucosal plexus by 22 weeks, mirroring human ENS development^60,61^. This experiment also highlighted the structural integrity and functionality of the engineered tissues in a fed intestinal state. By leveraging confinement to enhance cell-cell interactions and co-development, this methodology optimizes both in vitro and in vivo organoid maturation, paving the way for future research and therapeutic applications.

## ADDITIONAL INFORMATION

Supplementary information is available for this paper. Correspondence and requests for materials should be addressed to Maxime M. Mahe and/or Michael A. Helmrath.

## Supporting information

Poling_et_al_bioRxiv Supplementary File

## ACKNOWLEDGEMENTS

The authors thank Mya Michelle Bligen, Maksym Krutko and Emma Mayers for their support in completing animal work and Leyla Esfandiari for providing feedback on an early draft of the manuscript. Schematics were created with BioRender.com.

## FUNDING SOURCES

Research reported in this publication was supported by the National Institute of Diabetes and Digestive and Kidney Disorders (NIDDK) and the National Institute of Allergy and Infectious Diseases (NIAID) of the National Institutes of Health under grant number U01DK103117. The content is solely the responsibility of the authors and does not necessarily represent the official views of the National Institutes of H ealth. This work was also funded in part by NIH grant number NIH P30 DK078392, Cincinnati Digestive Health Center Award (Gene Expression Core, DNA Sequencing and Genotyping Core, Pluripotent Stem Cell Core). This work was supported in part by ANR-17-CE14-0021 (SyNEDI to M.M.M.), ANR-21-CE14-00 (NeuroPIMM to M.M.M.). This work was supported in part by a National Science Foundation GRFP Award (H.M.P.).

## AUTHOR CONTRIBUTIONS

HMP, MAH, and MMM conceived and designed the study. HMP designed scaffolding trays. MRB manufactured scaffolding tray molds. HMP, ALP, AS, KN, and NB performed cell culture. KT, PC and TN conducted study related bioinformatics. SM and IA generated RRG rat model for transplantation. HMP and MAH performed all surgical procedures. HMP, AS, GWF, KS, and TH performed assays, collected data, and/or performed analysis. HMP prepared figures and drafted the manuscript. HMP, RB, CNM, JMW, TT, MAH, and MMM made critical revisions to the manuscript, and all authors approved its final version.

## COMPETING INTERESTS

Cincinnati Children’s Hospital Medical Center has a patent in process associated with the methods established in this study.

## Methods

### Manufacturing Scaffolding Trays

Scaffolding tray molds were produced using a Form 2 3D printer (FormLabs) out of Surgical Guide Resin (FormLabs). Then, molds were filled with degassed polydimethylsiloxane and allowed to cure before removal and use.

### Human Tissue

Human tissue collection was performed with the approval of Cincinnati Children’s Hospital Medical Center’s (CCHMC) Institutional Review Board (Tissue Characterization, Study No. 2014-0427 and SUR IPSC and Tissue Engineering, Study No. 2017-2011). Surgical samples of pathologically normal human small intestine and colon were obtained from patients undergoing bariatric or revision/resection procedures between the ages of 14 and 25 years old to serve as control samples. Informed consent or assent was obtained from all patients and/or parent/legal guardians as appropriate. Additional deidentified samples of pathologically normal colon were obtained through CCHMC’s Discover Together Biobank. All human tissue was utilized in accordance with institutional ethics guidelines.

### Animals

All animal procedures and experiments were performed with the prior approval of Cincinnati Children’s Hospital Medical Center’s (CCHMC) Institutional Animal Care and Use Committee (Building and Rebuilding the Human Gut, Protocol No. 2021-0060 and No. 2024-0117). Male and female adult immunodeficient rats with Rag1 and Il2rg gene deletions (RRG) with ages between three and six months of age were used for conventionally generated and groove-based organoid transplantations (founders from Transgenesis Rat ImmunoPhenomic Platform, Nantes, France, in-house breeding)^17^. Additionally, adult immune-deficient NOD-SCID IL-2Rγ^null^ (NSG) mice with ages between three and six months of age were used in conventionally generated organoid experiments (Comprehensive Mouse and Cancer Core Facility, Cincinnati, Ohio). Animals were housed in CCHMC’s barrier animal vivarium and handled humanely in accordance with the NIH Guide for the Care and Use of Laboratory Animals.

### Human Pluripotent Stem Cell (hPSC) Lines

The H1 human embryonic stem cell line (WA01) was obtained from WiCell. The induced pluripotent stem cell line 72_3 was generated by the CCHMC Pluripotent Stem Cell Facility and has previously been described^6^. hPSCs were cultured under feeder-free conditions using mTESR1 (StemCell Technologies) and human embryonic stem cell-qualified Matrigel (Corning). Spontaneously differentiated cells were manually removed and media was changed daily.

### Organoid Generation

#### Small Intestinal Organoids

Aggregate and spontaneous spheroids were generated as previously described^8,14,62^. Briefly, hPSCs were grown in feeder-free conditions in Matrigel (BD Biosciences) coated six-well Nunclon surface plates (Nunc) and maintained in mTESR1 media (STEMCELL Technologies). For definitive endoderm (DE) induction, cells were passaged as single cell suspensions generated with Accutase (STEMCELL Technologies) and plated in 24-well Nunc plates at a density of approximately 100,000 cells/well. Cells grew in mTESR1 media for two days before treatment with 100 ng/ml of Activin A for three days. DE was then treated with hindgut induction medium (RPMI 1640, 100x NEAA, 2% dFCS,) for four days with 100 ng/ml FGF4 (R&D) and 3 µM Chiron 99021 (Tocris) to induce spontaneous free-floating mid-hindgut spheroids which would be collected for use. Additionally, the remnants of the underlying monolayer were used to generate aggregate mid-hindgut spheroids. After making a single cell suspension, cells were plated in Aggrewell 400 plates (StemCell Technologies) at approximately 3000 cells per microwell with intestinal growth medium (Advanced DMEM/F-12, N2 supplement, B27 supplement, 15 mM HEPES, 2 mM L-glutamine, penicillin-streptomycin) supplemented with 10 µM Y-27632. After 24 hours, aggregate spheroids were collected for use. Upon collection, spontaneous and aggregate spheroids were plated in Growth Factor Reduced (GFR) Matrigel and maintained in intestinal growth medium supplemented with 100 ng/ml EGF (R&D) to generate HIOs. Alternatively, for the Bailey Method to generate SI CCS, spheroids were plated within the grooves of the scaffolding trays with approximately 4,000 spheroids per groove in GFR Matrigel and maintained in intestinal growth medium. After six days, structures were removed from the grooves with forceps and re-plated as a longitudinal structure in untreated Tissue Train cell culture plates (Flexcell International) using the nylon tabs as anchors where culture continued until 14 days. Intestinal growth media was changed twice weekly for all organoids. HIOs were utilized for surgical transplantation on days 14 and 28. SI CCS were utilized for transplantation on day 14.

#### Small Intestinal Organoid Assembloid with Neural Crest Cells

Organoid assembloids with an ENS (HIO+ENS) were generated as previously described^26^. Briefly, mid-hindgut spheroids were combined with neural crest cells (NCCs), which were generated in parallel to the spheroids. NCCs were differentiated as previously described for neuroglial lineages and the spheroids were generated as described above^63^. To combine the spheroids and NCCs, a ratio of 1:600 was mixed and they were centrifuged for 3 minutes at 300g before embedding in Matrigel. The culture conditions that followed were identical to that of the conventional small intestinal organoids (HIOs) as outlined above.

#### Colonic Organoids

Spheroids were generated in an identical manner to those listed above in small intestinal organoid generation. To distalize the structures, BMP2 (100 ng/ml, R&D) was added to cultures upon plating spheroids, either conventionally or in the scaffolding trays, and changed to intestinal growth media after 72 hours^7^. HCOs were utilized for surgical transplantation on days 14 and 28. C CCS were utilized for transplantation on day 14.

#### Small Intestinal Organoids with Exogenous Neural Crest Cells

Spheroids were generated identically to those listed above in small intestinal organoid generation. In parallel to generating spheroids, neural crest cells were also differentiated and combined with spheroids as previously described^26^. Briefly, for neural crest cell (NCC) generation, hPSCs were treated with collagenase then gently broken up, and resuspended in neural induction media on non-tissue culture-treated petri dishes. Neural induction media was changed daily, and retinoic acid (2 µM) was added on days four and five f or p osteriorization. O n d ay s ix, f ree-floating ne urospheres we re plated on fibronectin (3 µg/cm^2^) and fed neural induction media for four d ays. Migrated cells were collected using an Accutase treatment and combined with spheroids at a ratio of 600:1 before replating in GFR Matrigel. HIO+ENS were utilized for surgical transplantation on day 28.

#### Stomach Organoids

Spontaneous anterior foregut spheroids were generated as previously described^5,6,11^. Briefly, hPSCs were grown in feeder-free conditions in Matrigel (BD Biosciences) coated six-well Nunclon surface plates (Nunc) and maintained in mTESR1 media (STEMCELL Technologies). For definitive endoderm (DE) induction, cells were passaged as single cell suspensions generated with Accutase (STEMCELL Technologies) and plated in 24-well Nunc plates at a density of approximately 340,000 cells/well. Cells were grown in mTESTR1 media for one day prior to differentiation. On the first day of differentiation, cells were treated with both 50 ng/mL BMP4 (R&D) and 100 ng/ml of Activin A (Cell Guidance Systems) in RPMI 1640 (Life Technologies). For the next two days, the cells were only treated with 100 ng/mL of Activin A in RPMI 1640. DE was then treated with foregut induction medium (RPMI 1640, 100x NEAA, 2% dFCS,) for three days with 500 ng/ml FGF4 (R&D), 2 µM Chiron 99021 (Tocris), and 200 ng/mL Noggin (R&D) to induce spontaneous free-floating mid-foregut spheroids which would be collected for use. 2 µM Retinoic Acid (RA) was added on the third day of FGF/Chiron/Noggin treatment. Spheroids were collected on Day 6 of differentiation and were then plated in Growth Factor Reduced (GFR) Matrigel and maintained in intestinal growth medium supplemented with 100 ng/ml EGF (R&D) to generate HaGOs. For the first three days after spheroid collection, intestinal growth medium was supplemented with 200 ng/mL Noggin and 2 µM RA. Alternatively, to generate G CCS, spheroids were plated within the grooves of the scaffolding trays with approximately 2,200 spheroids per groove in GFR Matrigel and maintained like their conventional counterpart. After nine days, structures were removed from the grooves with forceps and re-plated as a longitudinal structure in untreated Tissue Train cell culture plates (Flexcell International) using the nylon tabs as anchors where culture continued until 16 days. Intestinal growth media was changed twice weekly for all organoids. HaGOs were utilized for surgical transplantation on day 28. G CCS were utilized for transplantation on day 16.

#### Organoid Transplantation

Conventionally generated organoids, HIOs, HIO+ENS, HCOs, and HaGOs, and CCS organoids, SI CCS, C CCS and G CCS, were transplanted within the renal subcapsular space and within the mesentery as previously described^14–16^. Briefly, the kidney was accessed for kidney capsule transplantations, and a pocket was created under the kidney capsule for organoid insertion. For mesentery transplantations, the bowels were eviscerated from the thoracic cavity. A suitable bifurcating vessel was identified for the transplantation of an organoid using a single drop of topical tissue adhesive to secure the organoid (GLUture). Due to the increased organoid size, CCS organoids were only transplanted within the mesentery. Their placement was adjacent to the host’s distal small bowel or proximal colon instead of on a bifurcating vessel structure. Both food and water were provided ad libitum before and after surgeries. Doses of carprofen (5 mg/kg) and bupivacaine (1 mg/kg) were administered for pain management. Animals were monitored post-operatively and additional analgesics were administered as needed.

#### Tie In Surgical Procedure

Ten weeks following engraftment, mice and rats underwent a secondary surgery as described previously^39^. During this Tie In procedure, the engrafted organoid (tHIO or SI CCS) was incised using a surgical scalpel blade number 11 and luminal contents removed for collection. Then, the adjacent host bowel was incised, and luminal content removed using an 18G blunt tip fill needle and injected into the open lumen of the tHIO. Then, using 9-0 Ethilon suture (Ethicon) the tHIO and host bowel were anastomosed in a simple interrupted fashion. Cotton swabs were used to apply an inward rolling pressure along the joined structure to expose leaks and ensure the organoid was filled with luminal content of the host. In sham procedures, the organoid was incised, luminal contents removed before closing in a simple interrupted fashion with 9-0 Ethilon suture. Mice were sacrificed and tissue harvested 5d and 12d p ostoperatively. Rats were sacrificed and tissue harvested 12 weeks postoperatively.

#### Single Cell Extractions and Sequencing

Single cells were isolated from d28 HIO+ENS and d14 SI CCS in vitro structures. Organoids were carved out of their Matrigel and transferred to 1% BSA coated 15 mL conical tubes. Then, the organoids were incubated in Accutase (STEMCELL Technologies) containing Y-27632 (STEMCELL Technologies) for 10 minutes at room temperature with intermittent pipetting to mix. After 10 minutes, a small portion of the mixture was visually inspected using a microscope and an additional 2 minutes for digestion was allowed as needed. Then, any remaining fragments were allowed to settle and the liquid removed and added to a fresh 15 mL conical tube containing DMEMF12 with 10% FBS to stop further digestion. More Accutase containing Y-27632 was added to the remaining undigested organoid fragments and an additional 20 minutes allowed for digestion. Once ∼90% of the suspension was digested, DMEMF12 with 10% FBS was added to stop further digestion. Then both digestion fractions were pooled and filtered through a 40 µm cell s trainer. Cells were then washed, counted, and utilized for sequencing.

#### Single Cell RNA Sequencing Bioinformatics scRNA-seq Processing

Each sample was processed separately until the integration steps. FASTQ files are first processed and checked for quality using Cellranger (https://github.com/10XGenomics/cellranger), which aligns the reads in each sample against the human hg38 genome. The count matrix is then processing using Scanpy and a standard workflow. Doublets w ere estimated using the Scrublet tool^64^. Filtering then removed all genes expressed in less than 10 cells and all cells with less than 200 genes as well as cells expressing with than 5 percent of mitochondrial transcripts.

All count matrices were merged and then normalized following the shifted logarithm method. Batch aware 3000 highly variable genes were defined and used for downstream analysis. Data was scaled while regressing out unwanted variation from cell cycle phase and PCA was performed as a means of linear dimensionality reduction.

#### scRNA-seq Integration and Annotation

All samples were then integrated using Scanpy and the harmonypy package using the first 5 0 p rincipal components of the PCA^65^. A nearest neighbors approach was used to compute a distance matrix and a neighborhood graph of the cells which was subsequently used to compute a UMAP. Clusters were computed using the Leiden algorithm^66^ and differentially expressed genes between said clusters were identified using the sc.tl.rank_genes_groups() function of the Scanpy package. Based on these results and on the expression of a few selected markers, clusters were then identified as one of these populations of cells : Epithelial (*EPCAM, FABP1*), Mesenchymal (*COL1A2, COL3A1, MYH11, ACTG2, TAGLN*), Endothelial (*CD34, PROM1, KDR, PECAM1, ICAM1, CDH5, CLDN5*) and Neural (*B3GAT1, NGFR, NCAM1, PHOX2B, HAND2, TUBB2B, FOXD3, TFAP2A, ASCL1, ELAVL4, SOX10, S100B*).

#### scRNAseq maturation scores

Using lists of know marker genes, early (*SOX10, FOXD3, PHO2B, RET, EDNRB, PAX3, NES*) and late (*ELAVL3, ELAVL4, TUBB3, DCX, MAP2, NEFL, GAP43*) neuronal scores were computed for each cell in the dataset as the mean expression of respective marker genes.

#### scRNAseq similarity scores

To evaluate the maturity of identified neural cells, both the query and reference datasets -the Gut Cell Atlas’ subset of fetal neuronal cells were subsetted to include only common genes^66,67^. Mean expression profiles were calculated for each annotated cell type in the reference dataset. Similarity scores were then computed for each query cell against these reference profiles using Spearman correlation. The highest similarity score and its corresponding reference cell type annotation were identified and recorded for each query cell.

#### Single Nucleus Extractions and Sequencing

Nuclei were isolated from fresh frozen HIO, SI Groove, and Human Jejunum tissue samples using the Minute Detergent Free Nuclei Isolation Kit (Invent Biotechnologies, Inc.) following manufacturer guidelines. Immediately following completion, nuclei were assessed and submitted to CCHMC’s Gene Expression Core facility to complete 10X Genomics Chromium System assays using Chromium Single Cell 3’ Library & Gel Bead Kit v3 (10X Genomics). Then, sequencing was performed by CCHMC’s DNA Sequencing and Genotyping Core on a NovaSeq 6000 System (Illumina).

#### Single Nucleus RNA Sequencing Bioinformatics snRNA-seq Processing

FASTQ files are first processed and checked for quality using Cellranger (https://github.com/10XGenomics/cellranger), which aligns the reads in each sample against the human hg38 genome. SoupX is then run on the resulting output to correct for ambient RNA^68^. The corrected counts’ matrix is then processed using the standard Seurat workflow^69^. Filtering removed all genes expressed in less than 3 cells and all cells with less than 200 genes. Normalization uses a global scaling method to adjust and log-transform the expression values for each cell, and feature selection selects the 2000 genes with the highest cell-to-cell variation to be used in all downstream analyses. Data is also scaled to mean 0 and variance 1 while regressing out unwanted variation from mitochondrial RNA. PCA was performed as a means of linear dimensionality reduction, and cells are clustered with a k-nearest neighbors approach using the first 20 principal components and a resolution of X.UMAP was also performed to visualize these clusters, a type of nonlinear dimensionality reduction. Differentially expressed genes between clusters and cell types were identified with the ‘FindAllMarkers’ function, which uses the Wilcoxon rank-sum test by default.

#### snRNA-seq Integration

All samples were then integrated into a single dataset with Seurat, to remove batch effects and analyze them jointly. The method involves dimensionality reduction using diagonalized canonical correlation analysis (CCA), L2-normalization of the canonical correlation vectors (CCV), and finding mutual nearest neighbors (MNNs) in the low dimensional space between two datasets. These neighbors or “anchors” are compared and used to apply a correction vector to the dataset, which continues in a recursive pairwise manner between datasets until all are corrected. The first 30 canonical correlation components are used for clustering from this integrated dataset, which is then visualized with UMAP.

#### Constructing a single-cell reference atlas

To annotate the cell types present in the samples, a reference atlas was first constructed using the LabelHarmonization function in ELeFHAnt^70^. Three datasets from fetal gut development were used: E-MTAB-8901, GSE158702, and E-MTAB10187. Intestinal cells were subsetted and combined using the canonical correlation analysis-based integration in the R package Seurat. We also removed certain cell types that were not informative for our data, resulting in a final dataset of 70 cell types, 62 clusters, and ∼115,000 cells. This dataset was divided into training and test sets, and using an ensemble learning that combines predictions from multiple classifiers ELeFHAnt predicted cell types for each integration cluster. This harmonized the cell types to a final set of 41, that through manual curation had some cell types renamed based on marker analysis.

#### Annotation of snRNA-seq

The ClassifyCells_usingApproximation function in ELeFHAnt^70^ was used to predict cell types for the samples. The reference atlas served as the training set, samples were used as the test set, and the same ensemble learning approach mentioned previously was applied. This assigns the most frequently predicted reference cell type within each query cluster to that cluster. As an additional annotation method, the MapQuery function was used from Seurat. Following a similar method as seen with integration, similar pairs of cells or “anchors” are found between reference and query by projecting the reference PCA onto the query rather than using CCA. Using these anchors, a weighted vote classifier finds the highest confidence cell type in the reference to assign to each query cell. Finally, these predictions can be visualized by projecting the query onto the reference UMAP.

#### Bulk RNA Isolation and Sequencing

According to manufacturer guidelines, RNA was extracted from in vitro organoids using an RNeasy Plus Micro Kit (Qiagen). Then, samples were quantified and submitted to CCHMC’s DNA Sequencing and Genotyping Core for Next Generation Sequencing. All samples were assayed to have RNA integrity numbers greater than eight. After quality control, a cDNA library was created and sequenced using an IlluminaHiSeq2000 (Illumina) with 20 million paired-end reads per sample.

#### Bulk RNA Sequencing Processing

FASTQ files were processed with CSBB (https://github.com/csbbcompbio/CSBB-v3.0), which uses RSEM^71^ for mapping and quantification. Paired-end reads were aligned against the human reference genome hg38 using bowtie2. All samples were checked for quality using FastQC metrics (https://www.bioinformatics.babraham.ac.uk/projects/fastqc/).

#### Bulk RNA Sequencing Analysis

The count matrices generated were first filtered to remove long non-coding RNAs. DESeq2^72^ was then used to output differentially expressed genes by comparing genes between conditions of interest. For PCA, the normalized counts matrix (in transcripts per million) is log-transformed and principal components are calculated. Figures for the PCA loadings are produced using the R package pcaExplorer^73^. Heatmap of differentially expressed genes were generated using logtransformed counts using R package pheatmap (https://cran.r-project.org/web/packages/pheatmap/index.html).

#### Tissue Processing and Immunostaining

Tissue samples were handled and processed as previously described^12^. Briefly, 4% paraformaldehyde (PFA) was used for overnight fixation at 4 °C and tissues were processed in a HistoCore Pearl (Leica) on a 12-hour cycle and embedded in paraffin blocks. Sections were deparaffinized and rehydrated to water and either stained immediately with hematoxylin and eosin or subjected to heat induce epitope retrieval, and antibody stained. 10% normal serums were used as blocking buffers. Primary antibody incubations took place at 4 °C overnight in 1% bovine serum albumin in phosphate-buffered saline (PBS). Antibodies and their respective dilutions are listed in Table S1. The Vectastain ABC system was used for amplification and the diaminobenzidine substrate kit was used for signal detection (Vector Laboratories). Lillie-Mayer’s Hematoxylin (Agilent Technologies) was used as a counterstain.

#### Image Acquisition

Surgical imagery was acquired using an M80 microscope outfitted with a MC 170HD camera (Leica Microsystems). Gross images of harvested structures were acquired using a V60 ThinQ (LG Electronics, Inc.). Harvests were performed using a M165 FC microscope outfitted with a DCF7000 T camera (Leica Microsystems). Slides were imaged using an Eclipse Ti microscope (Nikon Corporation) and subsequent analysis was performed using Nikon Elements Imaging Software (Nikon Corporation).

#### Organoid Size and Morphometric Analysis

Engrafted organoids were photographed with a ruler and quantified using ImageJ (National Institutes of Health) with a conversion factor of 1 mm as measured on the ruler. Morphometric analysis was performed on hematoxylin and eosin-stained tissue sections. Crypt depth, crypt width, villus height, villus width, and mucosal thickness were measured for a minimum of 20 well-oriented crypt-villus units per tissue sample and then averaged using Nikon NIS imaging software (Nikon). Specific regions or image frames were not used, this was performed across the entire tissue contained on each slide and each data point represents the average from a biological replicate. For neuronal bundle size quantifications, cross sectional areas were measured in Nikon NIS imaging software (Nikon). Again, this was not from a single image frame, but the entirety of the section.

#### Ex vivo Neuromuscular Function and Muscle Tone

Organ bath chamber assays were performed as previously described^11,12,26^. The muscularis of freshly harvested organoid samples were carefully dissected into small strips of approximately 2 mm x 6 mm; no chelation buffer was used. Muscle strips were mounted within an organ bath chamber system for assessment (Radnoti) to force transducers (ADInstruments) and contractile activity was continuously recorded using LabChart software (ADInstruments). After an equilibration period, a logarithmic dose response to Carbamyl-β-methylcholine chloride (Bethanechol; Sigma-Aldrich) was obtained by administering exponential doses with concentrations of 1 nM to 10 mM at 2 min intervals before the administration of 10 µM scopolamine (Tocris Bioscience). After another equilibrium period, muscle preparations were then stimulated with a control electrical field stimulation (EFS) pulse. The EFS protocol was programmed as follows: 8000 mV pulse for 400 µs followed by -1000 mV pulse for 400 µs and a 400 µs rest for a total of 10 sec. NG-nitro-L-arginine methyl ester (L-NAME; 50 µM; Sigma) was applied 10 min before EFS stimulation to observe the effects of NOS inhibition. Without washing, Atropine (atropine sulfate salt monohydrate; 1 µM; Sigma) was applied 10 min prior to another EFS stimulation to observe the cumulative effect of NOS and Ach receptor inhibition. After several washes and an additional equilibrium period, another control EFS pulse was administered. Neurotoxin tetrodotoxin (TTX; 4 µM; Tocris) was administered 5 min before a final EFS stimulation. Analysis was performed by calculating the integral (expressed as the area under the curve, AUC) immediately before and after stimulation for 60 s. For cholinergic subtype calculations, the change is response associated with L-NAME exposure was subtracted from the total response and divided by the total response. Then, the change in response associated with both L-NAME and Atropine exposures was subtracted from the total response and divided by the total response to determine the other types. Finally, both values were subtracted from the whole to determine the nitrergic proportion. All data are normalized to muscle strip mass.

#### Ex vivo Epithelial Characteristics and Permeability

The epitheliums of freshly harvested sham and Tie In grafts were carefully dissected through a technique similar to seromuscular stripping as previously reported^12,74,75^. Samples were incised and dissection was done in ice cold Kreb’s buffer (NaCl, 117 mM; KCl, 4.7 mM; MgCl2, 1.2mM; NaH2PO4, 1.2 mM; NaHCO3, 25 mM; CaCl2, 2.5 mM and glucose, 11 mM). Full thickness tissue segments were then pinned in a dish containing 0.5 cm thick cured Sylgard (Electron Microscopy Sciences). The seromusculature was then micro-dissected as one unit from the epithelium using Dumont #5 and #7 forceps along with Vannas scissors (Fine Science Tools, Inc.). Gross tissue integrity was assessed using the stereoscope’s bottom lighting for uniformity in appearance and any damaged areas were removed. Some remnant subepithelial mucosa remained after dissection, as previously described^12^. The central portion of the epithelium, with the least amount of handling, was then carefully positioned for mounting between the hemi-chambers of an Ussing apparatus (Physiologic Instruments). 0.008 cm^2^ of tissue was exposed to 3 mL of oxygenated Krebs buffer at 37 °C throughout the assay. The transepithelial potential difference was detected with two paired electrodes affixed to a salt bridge containing 3% agar in 3 M KCl. The electrodes were connected to a VVC MC8 voltage clamp amplifier (Physiologic Instruments, San Diego). Electrode potential difference and fluid resistance values were offset to zero immediately before sliders were mounted between the chambers. A 30 min monitoring period was allowed for the establishment of equilibrium. Then, tissues were voltage-clamped at 0 mV while continuously measuring the short circuit current (Isc). For FITC-dextran permeability, 2.2 mg/ml FITC-dextran was added into apical side, and a sample was taken from the basolateral side every 30 minutes for 3 hours, replacing the same amount of fresh modified Kreb’s buffer in the basolateral side. Once all aqueous samples were collected, they were quantified with a plate-reader (Synergy 2, BioTek).

#### Data Representation, Statistics and Reproducibility

For bar graphs, data is represented as a mean ± standard deviation, with all individual data points represented. A dashed line indicates the mean and interquartile ranges for violin plots, representing all individual data points. Wilcoxon SignedRanks tests were performed for statistics of two groups of paired data. For statistics comparing unpaired data a student t-test was performed if n<5, for *n ≥* 5 a Mann Whitney was used. The statistical significance cutoff was p < 0.05 and confidence was 95%.

#### Data Availability

The authors declare that all data supporting the findings of this study are available within the paper and its supplementary information. Sequencing data have been made publicly available on the European Nucleotide Archive (ENA) under project accession PRJEB84049.

## Notes

### Competing Interest Statement

Cincinnati Childrens Hospital Medical Center has a patent in process associated with the methods established in this study.

