## Supplementary material for "Engineering Large-Scale and Innervated Functional Human Gut for Transplantation": Poling_et_al_bioRxiv Supplementary File

### Supplemental Information

### Table of Contents

Supplemental Figure 1……………………………………………………………….…2
**Scaffolding tray design and in vitro SI CCS growth.**

Supplemental Figure 2……………………………………………………………….…3
**Intestinal morphology is enhanced in SI CCS.**

Supplemental Figure 3……………………………………………………………….…5
**Neural cell populations are well represented in SI CCS.**

Supplemental Figure 4……………………………………………………………….…6
**Neurogenesis is enriched in SI CCS prior to transplantation.**

Supplemental Figure 5……………………………………………………………….…7
**C CCS and G CCS recapitulates functional human colonic and gastric tissues.**

Supplemental Figure 6……………………………………………………………….…9
**Modeling the short-term fed intestinal state can be done with HIOs in mice but is associated with high mortality.**

Supplemental Table 1……………………….……………………………………………11
**List of antibodies used for immunostaining.**

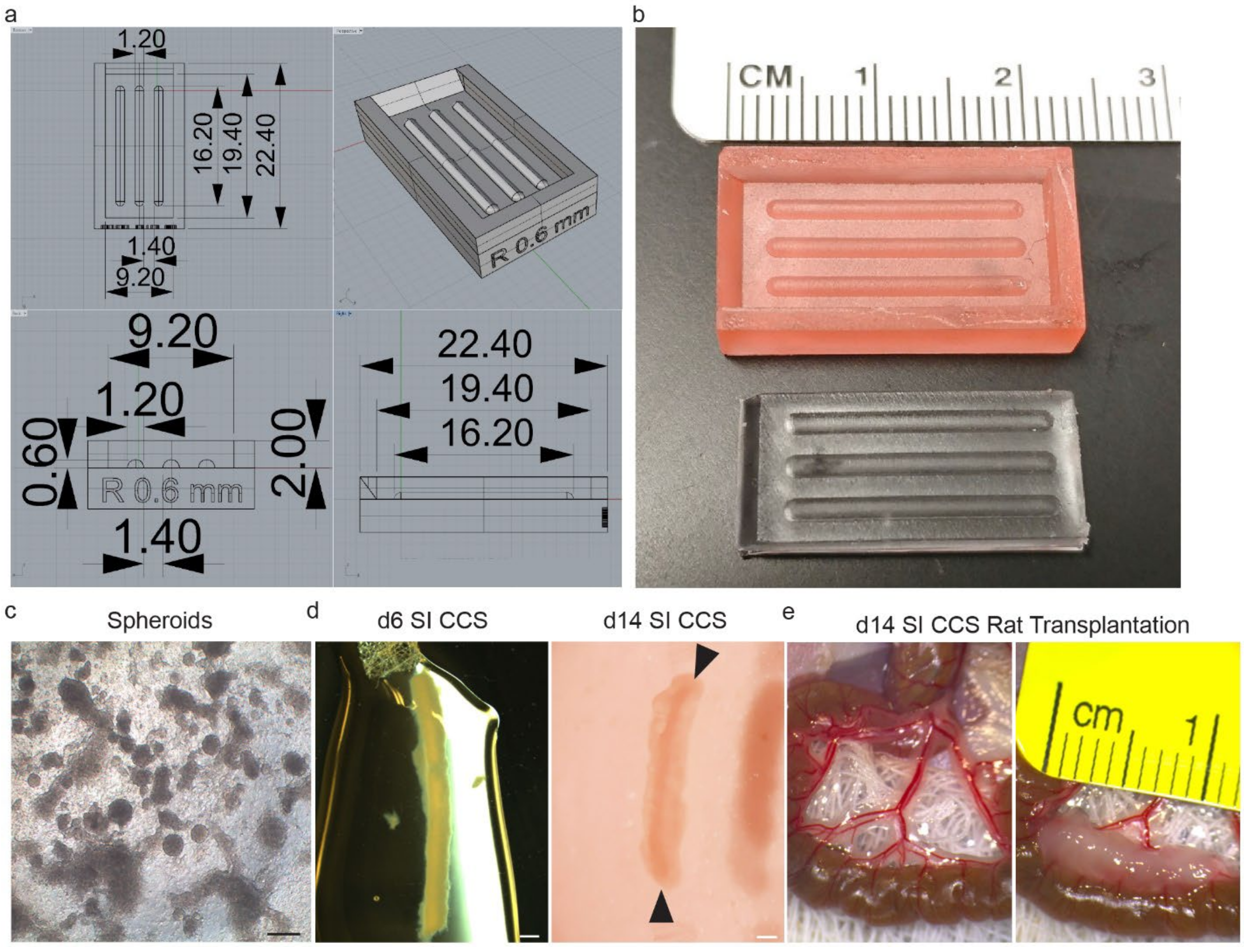


**Supplemental Figure 1. Scaffolding tray design and in vitro SI CCS growth.**
**a**, Schematic of scaffolding tray mold. **b**, Photograph of 3D printed scaffolding tray mold and PDMS scaffolding tray. **c**, Images of spheroids generated by spontaneous morphogenesis. Scale bars = 100 µm. **d**, Representative image of freshly replated d6 SI CCS after removal from scaffolding tray (left) and d14 SI CCS (right). Arrows indicate ends of dark longitudinal luminal area. Scale bars = 1 mm. **e**, Photographs of d14 SI CCS transplantation. **f**, Photograph demonstrating the central lumen after 10 weeks of engraftment.

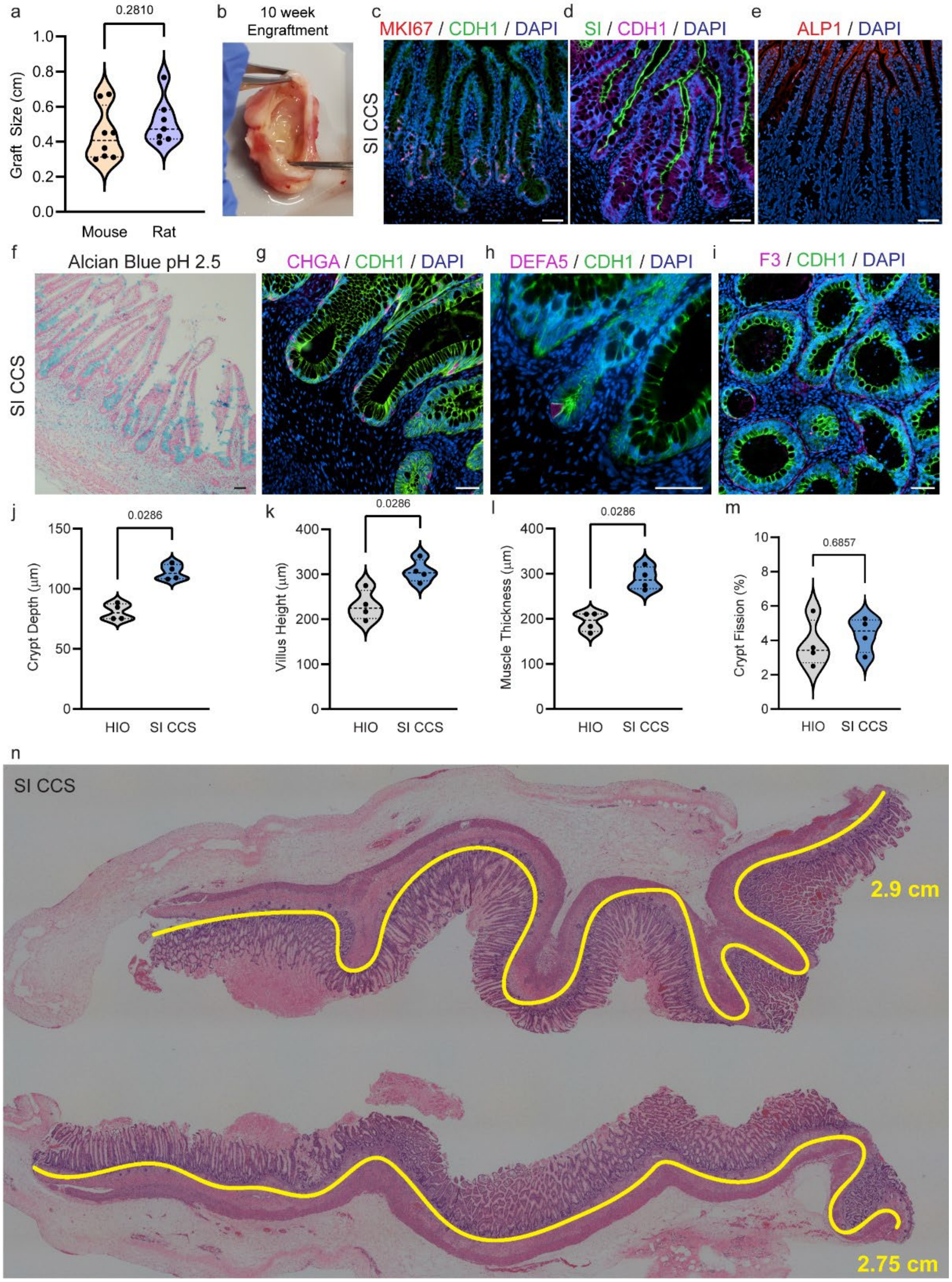


**Supplementary Figure 2. Intestinal morphology is enhanced in SI CCS.**

**a**, Violin plot of transplanted HIO size with mouse and rat as hosts. **b**, Photograph demonstrating the SI CCS central lumen after 10 weeks of engraftment. **c,** Representative staining for proliferation (MKI67, red) and epithelium (CDH1, green) in SI CCS. **d**, Representative staining for a brush border enzyme (SI, red) and epithelium (CDH1, green) in SI CCS. **e**, Representative image of active alkaline phosphatase activity in (ALP1, red) in SI CCS. **f**, Representative image of Alcian Blue pH 2.5 staining for mucins (blue) in SI CCS. **g**, Representative staining for enteroendocrine cells (CHGA, red) and epithelium (CDH1, green) in SI CCS. **h**, Representative staining for Paneth cells (DEFA5, red) and epithelium (CDH1, green) in SI CCS. **i**, Representative staining for telocytes (F3, red) and epithelium (CDH1, green) in SI CCS. **j-m**, Violin plots of morphometric quantifications of crypt depth (j), villus height (k), muscle thickness (l), and crypt fission (m) in transplanted HIOs and SI CCS. **n**, Tile scan of hematoxylin and eosin stained sections of SI CCS with yellow line highlighting continuous lengths of epithelium in sample. n=3 for all staining panels. Scale bars in **c-i** = 50 μm.

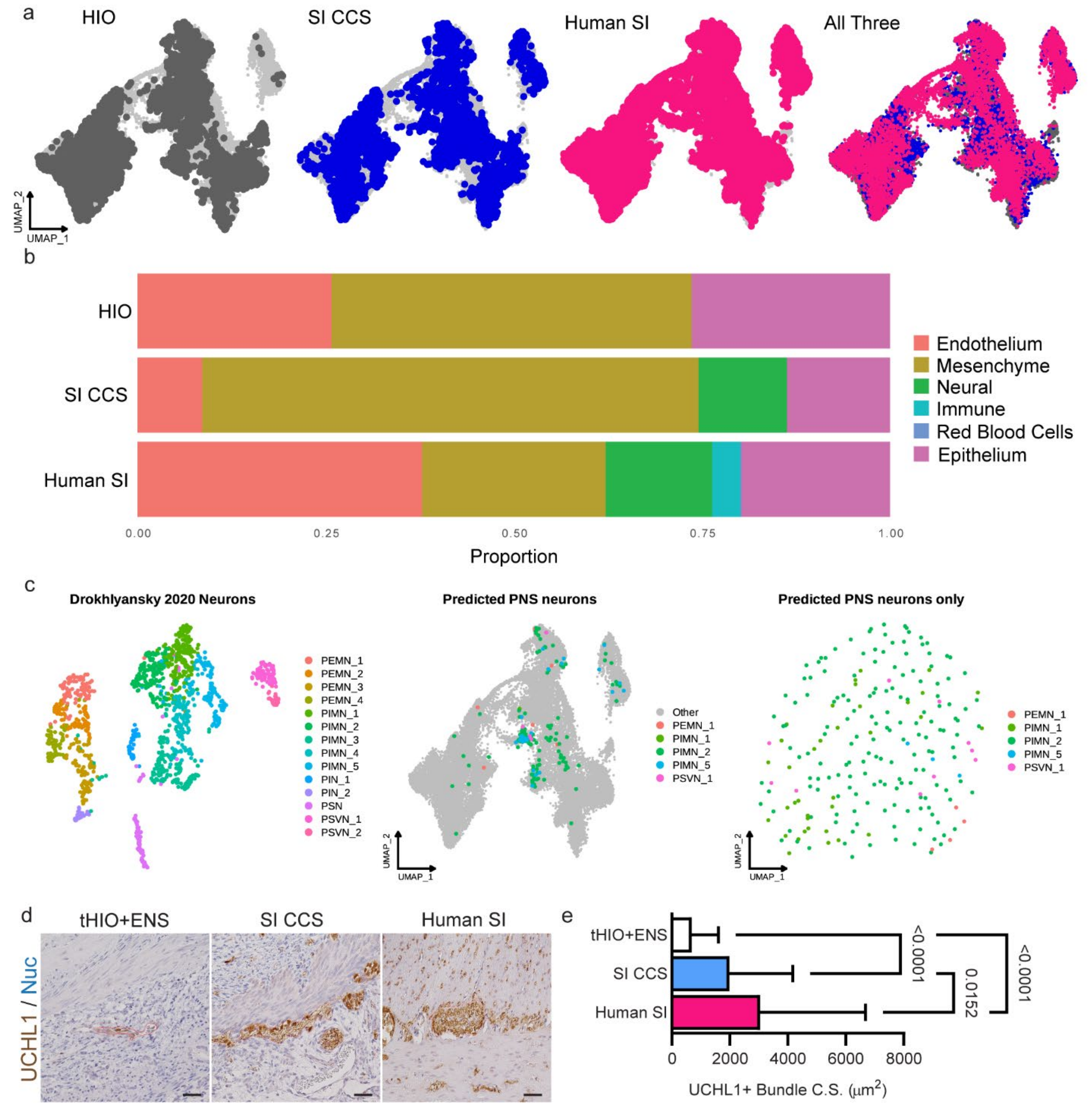


**Supplemental Figure 3. Neural cell populations are well represented in SI CCS.**
**a**, UMAPs of transplanted HIO, SI CCS and Human SI data sets projected individually onto the reference and combined. **b**, Bar graph representation of cell type proportions in transplanted HIO, SI CCS and Human SI. **c**, UMAPs of SI CCS neural cells projected neuronal classes using the Drokhlyansky dataset as reference. **d**, Representative staining for neurons (UCHL1, brown) in transplanted HIO+ENS, SI Grooves, and human small intestine. Scale bars = 100 µm. **e**, Bar graph (mean+SD) of measured cross sectional UCHL1+ bundle sizes from samples in panel d. Bundle sizes in SI CCS were significantly larger than those in transplanted HIO+ENS. n=3 per group.

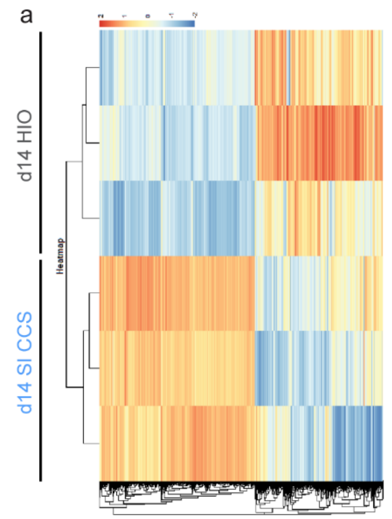


b

| Ontology | Biological Process | P Value |
|---|---|---|
| GO:0022008 | neurogenesis | 2.59E-44 |
| GO:0048699 | generation of neurons | 7.02E-43 |
| GO:0030182 | neuron differentiation | 7.89E-42 |
| GO:0007471 | central nervous system development | 2.07E-33 |
| GO:0007049 | cell cycle | 3.14E-33 |

| Ontology | Cellular Component | P Value |
|---|---|---|
| GO:0030424 | axon | 1.44E-25 |
| GO:0045202 | synapse | 7.26E-24 |
| GO:0043005 | neuron projection | 3.15E-23 |
| GO:0097060 | synaptic membrane | 8.56E-22 |
| GO:0098794 | post synapse | 9.10E-22 |

c

Events Captured in Single Cell Transcriptomics

| Organoid | Cell Type | Number |
|---|---|---|
| HIO+ENS | endothelial | 3 |
| | epithelial | 1,666 |
| | mesenchymal | 4,152 |
| | neural | 3,193 |
| | total | 9,014 |
| SI CCS | endothelial | 69 |
| | epithelial | 2,612 |
| | mesenchymal | 33,869 |
| | neural | 1,352 |
| | total | 37,902 |

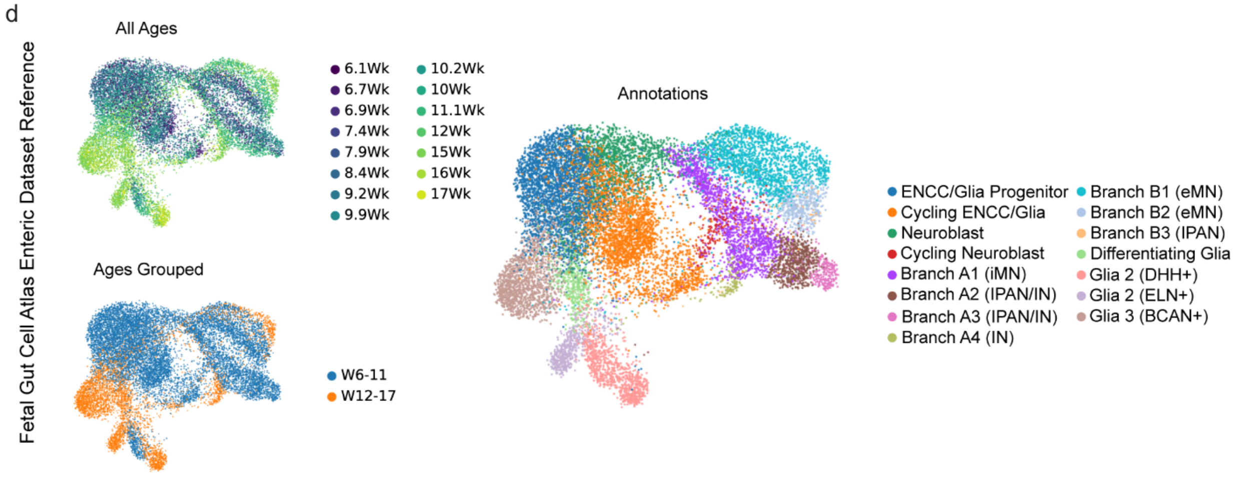


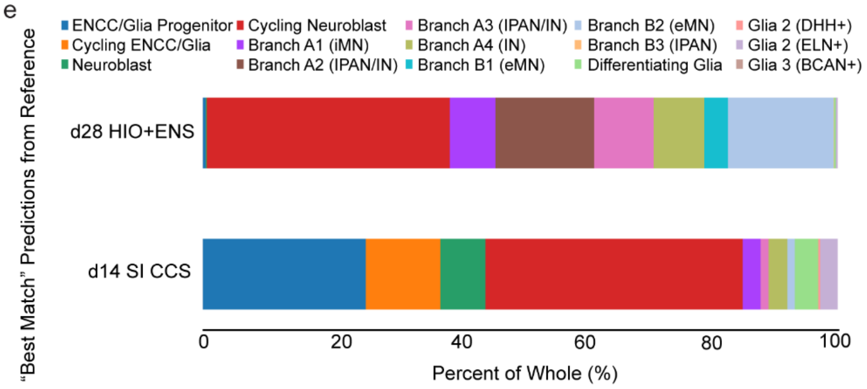


| | d28 HIO+ENS | d14 SI CCS |
|---|---|---|
| ENCC/Glia Progenitor | 14 | 347 |
| Cycling ENCC/Glia | 1 | 159 |
| Neuroblast | 6 | 96 |
| Cycling Neuroblast | 1221 | 548 |
| Branch A1 (iMN) | 231 | 38 |
| Branch A2 (IPAN/IN) | 496 | 0 |
| Branch A3 (IPAN/IN) | 299 | 17 |
| Branch A4 (IN) | 254 | 40 |
| Branch B1 (eMN) | 119 | 0 |
| Branch B2 (eMN) | 532 | 16 |
| Branch B3 (IPAN) | 2 | 0 |
| Differentiating glia | 9 | 50 |
| Glia 1 (DHH+) | 0 | 4 |
| Glia 2 (ELN+) | 9 | 37 |
| Glia 3 (BCAN+) | 0 | 0 |

**Supplementary Figure 4. Neurogenesis is enriched in SI CCS prior to transplantation.** **a**, Heatmap of bulkRNAseq data in d14 HIO and d14 SI CCS. **b**, Un-curated list of enriched gene ontologies in d14 SI CCS compared to d14 HIO. **c**, Table of cell type designations for scRNAseq analysis of d28 HIO+ENS and d14 SI CCS and their corresponding event (cell) counts. **d**, Fetal Gut Cell Atlas enteric reference curation into two time frames (early, w6-11, and late, w12-17) and associated cell type annotations. **e**, Bar graph and table of "Best Match" neural cell type predictions from the curated Gut Cell Atlas reference for d28 HIO+ENS and d14 SI CCS.

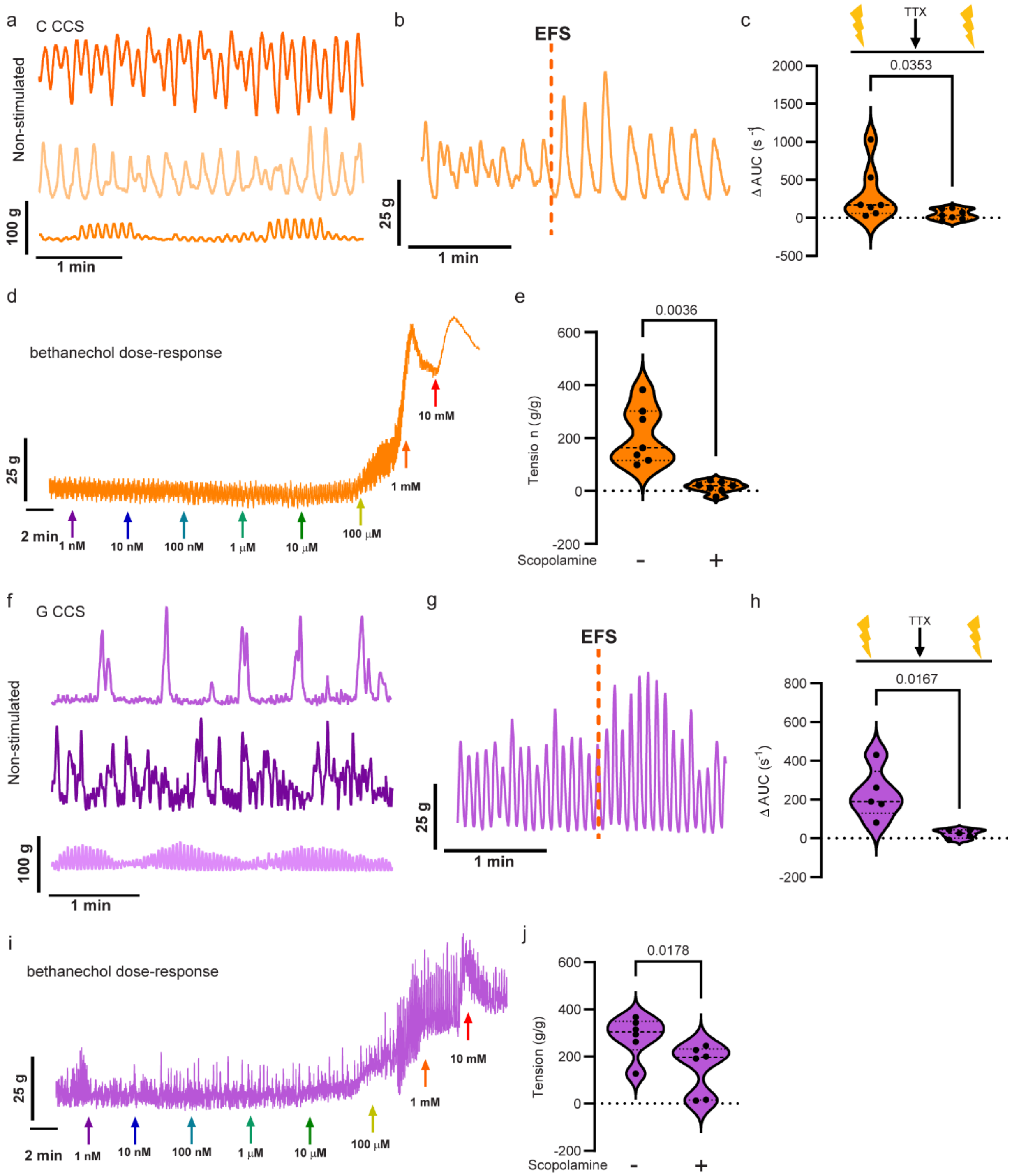


**Supplementary Figure 5. C CCS and G CCS recapitulates functional human colonic and gastric tissues.**

**a**, Contractile activity tracings in C CCS under homeostatic conditions. **b**, Contractile tracing in C CCS with EFS stimulation (dashed line indicates time of administration). **c**, TTX inhibition of ENS activity in C CCS. Violin plot of the change in the area under the curve before and after TTX. **d**, Representative bethanechol dose-response tracing of contractile activity in a transplanted C CCS. Colored arrows indicate the timing of logarithmic bethanechol doses. **e**, Calculated maximal and minimal tissue tension of C Groove; scopolamine was used to induce muscle relaxation. **f**, Contractile activity tracings in G CCS under homeostatic conditions. **g**, Contractile tracing in G CCS with EFS stimulation (dashed line indicates time of administration). **h**, TTX inhibition of ENS activity in G CCS. Violin plot of the change in the area under the curve before and after TTX. **i**, Representative bethanechol dose-response tracing of contractile activity in a transplanted G CCS. Colored arrows indicate the timing of logarithmic bethanechol doses. **j**, Calculated maximal and minimal tissue tension of G Groove; scopolamine was used to induce muscle relaxation. n=7 C CCS and n =6 G CCS for organ bath experiments.

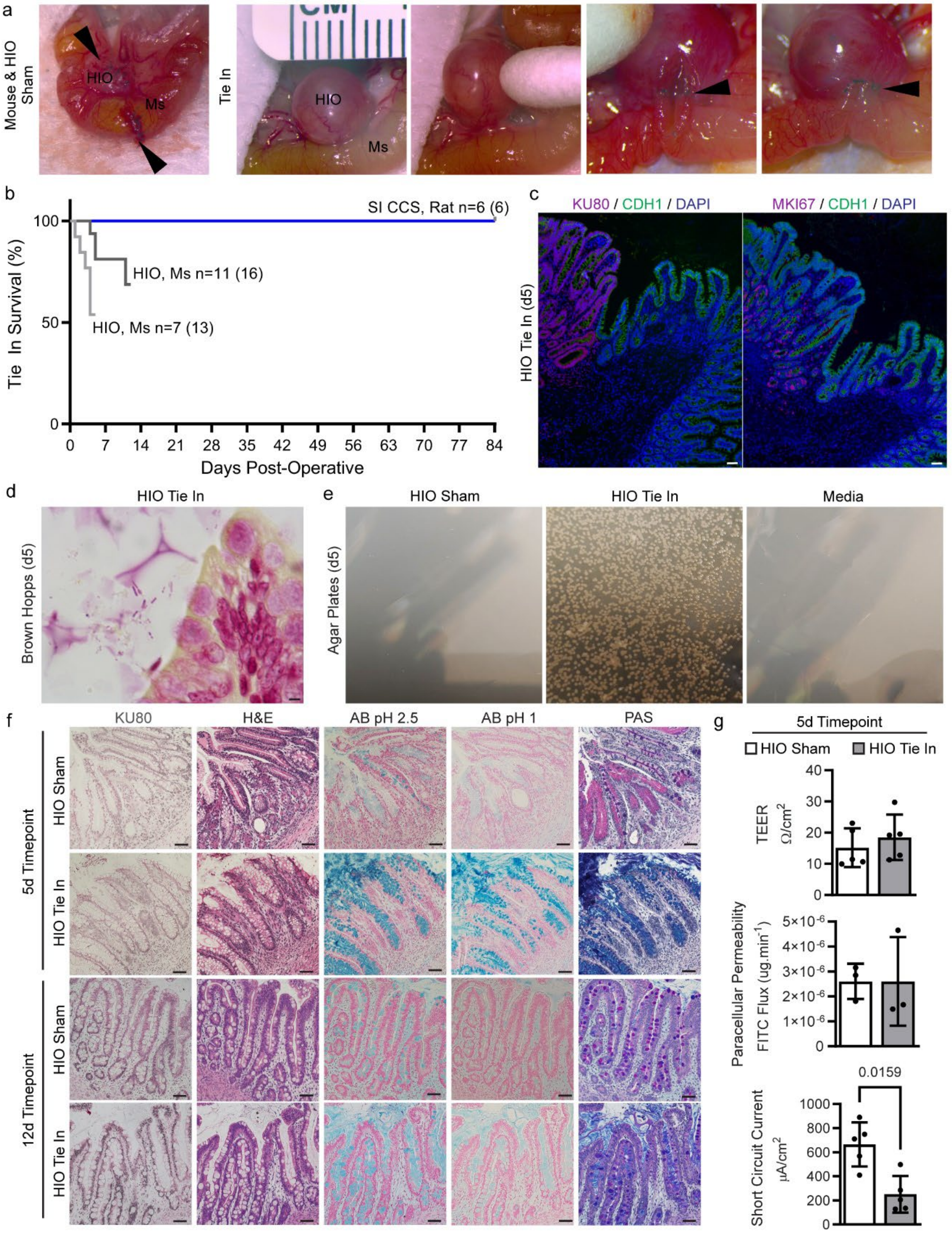


**Supplementary Figure 6. Modeling the short-term fed intestinal state can be done with HIOs in mice, but is associated with high mortality.**

**a**, Surgical images of HIO Sham and Tie In procedures. Sites of anastomosis are marked with black arrowheads. HIO and mouse tissues are denoted with text. **b**, Kaplan-Meier curve of survival associated with 5 day HIO Tie Ins, 12 day HIO Tie Ins and 12 week SI CCS Tie Ins. c,

Representative immunofluorescence staining for human cells (KU80, magenta) and epithelium (CDH1, green) demonstrating continuity between organoid and host (left). Serial section stained for proliferation (MKI67, magenta) and epithelium (green) demonstrating proliferative zonation within the intestinal epithelium. Scale bars = 50 μm. **d**, Representative Brown-Hopps staining for bacteria in HIO Tie In demonstrating bacteria near the epithelium. Scale bars = 5 μm. **e**, Photographs of agar plates after 72 hours of culture using luminal contents from HIO Sham, HIO Tie In, and media alone as a negative control. Colony outgrowth was only visible in HIO Tie In samples. **f**, Representative staining panels of mucin profiles across HIO Sham and Tie In at the 5d and 12d timepoints. A human marker (KU80, gray) was used to validate tissue source in serial sectioned panels of histological stains (left to right: H&E, alcian blue pH 2.5, alcian blue pH 1.0, and Periodic acid-Schiff). Scale bars = 50 μm. **g**, Bar graphs (mean±SD) of functional epithelial characteristics in HIO Sham and HIO Tie In at d5 as measured in an Ussing Chamber assay. From top to bottom: transepithelial resistance, paracellular permeability and short circuit current. n=3 per group for staining and n = 5 per group for electrophysiology measurements.

| | Antigen | Dilution | Host | Company: Catalog Number |
|---|---|---|---|---|
| **Primary** | ATP4B | 1:500 | mouse | Invitrogen: MA3-923 |
| | CA1 | 1:800 | rat | R&D: MAB2180 |
| | CD31 | 1:200 | rabbit | Abcam: ab28364 |
| | CDH1 | 1:500 | mouse | BD: 610182 |
| | CHAT | 1:200 | rabbit | Millipore: AB-143 |
| | CHGA | 1:500 | mouse | DSHB: CPTC-CHGA-1 |
| | CLDN18 | 1:800 | rabbit | Atlas: HPA018446 |
| | DEFA5 | 1:500 | mouse | Abcam: ab90802 |
| | F3 | 1:200 | rabbit | Atlas: HPA049292 |
| | GHRL | 1:500 | goat | Novus: NB600-813 |
| | HuC / HuD | 1:200 | mouse | Invitrogen: A21271 |
| | KIT | 1:300 | rabbit | Abcam: ab32363 |
| | MKI67 | 1:500 | rabbit | Thermo: RM-9106-S0 |
| | MUC5B | 1:900 | rabbit | Atlas: HPA008246 |
| | MUC6 | 1:1000 | mouse | Novus: NBP2-47799 |
| | NES | 1:400 | mouse | Abcam: ab22035 |
| | NF200 | 1:2000 | chicken | Antibodies-Online: ABIN953663 |
| | NOS1 | 1:200 | rabbit | Abcam: ab76067 |
| | S100B | 1:300 | rabbit | Atlas: AMAb91038 |
| | SATB2 | 1:200 | rabbit | Millipore Sigma: 384R-14 |
| | SI | 1:1000 | rabbit | Thermo: HPA011897 |
| | SOX2 | 1:500 | rabbit | GeneTex: GTX101507 |
| | SYN1 | 1:300 | rabbit | Abcam: ab8 |
| | TUBB3 | 1:350 | chicken | Abcam: ab41489 |
| | UCHL1 / PGP9.5 | 1:750 | rabbit | Dako: Z511601 |
| | XRCC5 / KU80 | 1:350 | rabbit | Cell Signaling: 2180 |
| **Secondary** | anti-chicken AF555 | 1:1000 | donkey | Thermo: A21449 |
| | anti-goat AF647 | 1:1000 | donkey | Thermo: A21447 |
| | anti-mouse AF555 | 1:1000 | donkey | Thermo: A21137 |
| | anti-rabbit AF488 | 1:1000 | donkey | Thermo: A21206 |
| | anti-rat AF488 | 1:1000 | donkey | Thermo: A21208 |

**Supplemental Table 1. List of antibodies used for immunostaining.**
